# MICAFlow: Fast and robust MRI preprocessing bridging research neuroimaging and clinical practice

**DOI:** 10.64898/2026.05.26.727725

**Authors:** Ian Goodall-Halliwell, Jordan DeKraker, Paul Bautin, Daniel Mendelson, Donna Gift Cabalo, Ella Sahlas, Alexander Ngo, Ke Xie, Jack Lam, Meaghan Smith, Youngeun Hwang, Laura Vavassori, Pietro Milano, Judy Chen, Arielle Dascal, Rui Ding, Guan Zhou, Marlo Naish, JiaJie Mo, Fatemeh Fadaie, Raul R Cruces, Boris C. Bernhardt

## Abstract

MICAFlow is a fully automated MRI preprocessing pipeline designed to translate advanced neuroimaging workflows from research into routine clinical practice. The pipeline emphasizes speed, robustness, and ease of use, focusing on structural and diffusion MRI. Key innovations include a Label-Augmented Modality-Agnostic Registration (LAMAReg) technique driven by deep learning segmentations for reliable cross-modal alignment, integration of state-of-the-art distortion corrections, and adherence to reproducible standards (Snakemake workflow, BIDSApp specifications). We describe the design of MICAFlow and evaluate its performance across heterogeneous datasets. First, accessibility: MICAFlow processes a multimodal MRI exam in minutes with clinically accessible hardware and without requiring GPU access, making it feasible for same-day clinical use. Second, registration accuracy: LAMAReg achieves cutting-edge multi-modal registration accuracy, yielding accurate alignment of diffusion MRI, FLAIR, and intra-subject T1-weighted images while remaining generally robust to common artifacts. Third, data reliability: Using identifiability, we show MICAFlow maintains consistent performance across diverse datasets, including subjects with pathology, and is closely comparable to contemporary pipelines. In sum, MICAFlow’s combination of machine learning and efficient workflows produces research-grade data quality with clinical-grade speed. This work demonstrates that advanced MRI preprocessing can be done fast and robustly, helping close the gap between research neuroimaging and broad clinical application of quantitative MRI techniques. The source code for MICAFlow is available here: https://github.com/MICA-MNI/micaflow, and for LAMAReg here: https://github.com/MICA-MNI/LAMAReg.

## Introduction

Despite advances in quantitative analysis of MRI data, routine clinical image interpretation remains predominantly visual, with quantitative tools still unevenly integrated into practice^1, 2^. Some clinicians have also expressed skepticism about the integration of advanced imaging approaches into clinical radiology, owing to concerns about transparency, limited validation and generalizability, and uncertainty about real-world workflow benefit^2–5^. Although a growing ecosystem of open neuroimaging pipelines supports quantitative MRI data analysis^6–9^, routine clinical image viewing and basic post-processing are still commonly embedded in vendor-managed radiology workflows, which can limit interoperability and local deployment of tools^6^. Additionally, many recent neuroimaging pipelines prioritize surface-derived outputs^8, 10, 11^ and/or achieve their fastest performance with GPU-accelerated components^8–10^, whereas surface-based biomarkers are not yet widely incorporated into routine clinical practice and computational infrastructure remains a recognized barrier to clinical deployment^4^. As a result, these tools can be poorly aligned with common clinical neuroradiology workflows, which often require rapid interpretation of multi-sequence volumetric brain MRI (for example, T1- or T2-weighted/Fluid Attenuated Inversion Recovery (FLAIR)-, and diffusion MRI-based protocols) in emergency or inpatient settings and operate under explicit report turnaround-time expectations^12^. Thus, beyond algorithmic performance alone, successful clinical deployment requires MRI analysis pipelines that are interoperable, accessible, and compatible with time-constrained neuroradiology practice.

A critical barrier to clinical adoption of MRI processing pipelines is the computational and technical bottleneck. Of note is the challenge of diffusion MRI (dMRI) processing, which typically requires time-intensive steps to correct for motion, eddy currents, and susceptibility distortions. Performing these steps on current pipelines can take hours, even on high-performance computing clusters^13, 14^. While these time demands are more affordable for research, clinically these delays can be meaningful^15^. The time-intensive and high-quality reconstructions are essential for more complex tractographic analyses, where even modest susceptibility-related geometric distortions can substantially alter reconstructed fiber pathways and connectivity estimates^16, 17^. In contrast, current clinical practice often relies on simpler diffusion-derived scalar metrics such as apparent diffusion coefficient (ADC) and fractional anisotropy (FA), which have shown good reproducibility and relatively low variability across preprocessing pipelines and acquisition settings^18, 19^. ADC represents the overall magnitude of water mobility within a voxel, while FA reflects the degree of directional dependence of water diffusion^20, 21^. A European-wide survey found that ADC maps were overwhelmingly used qualitatively (78% of respondents), by visual inspection only^22^. In addition, recent work has found that dMRI-derived metrics could aid detection of microstructural injury in traumatic brain injury^23^, help distinguish tumor infiltration from edema^24^, and have sensitivity in the assessments of pharmaco-resistant epilepsy^25^; however, broader clinical translation of quantitative dMRI has been slowed by the need for extensive artifact correction, rigorous quality control, and standardized, reproducible processing workflows^26–29^.

A further challenge is multi-modal image registration. Clinical MRI protocols routinely acquire multiple contrasts, notably T1-weighted (T1w), FLAIR or T2-weighted (T2w) MRI, and dMRI, with the exact sequence set tailored to the diagnostic question^30^. For accurate visual analysis, these images need to be co-registered within a common space. Many neuroimaging tools prioritize registration to a standard template for cross-subject analysis; in routine clinical interpretation, however, images are usually reviewed in native space, and spatial normalization becomes less straightforward in the presence of focal lesions, mass effect, or surgical cavities^31^. A key issue is that aligning images of different modalities is difficult because their intensity patterns differ and can be distorted by modality-specific artifacts. Traditional intensity-based or information-theoretic registration methods, such as mutual-information-based approaches, often struggle when modalities differ markedly in contrast or are affected by modality-specific artifacts, as in dMRI→T1w registration with echo-planar imaging (EPI) distortion or in clinical FLAIR→T1w registration with different contrast, resolution, and slice spacing^32^. Misregistration can impair diagnosis and lesion localization across sequences and can affect downstream clinical tasks that require millimetric spatial accuracy^33, 34^. Software tools often rely on boundary-based methods for robust cross-modal alignment, such as FSL’s epi_reg^13^ and FreeSurfer’s bbregister^35^. However, these approaches depend on having a clear and intact gray-white matter boundary, which may be less reliable when boundaries are obscured or displaced by lesional tissue. Despite many available registration algorithms, robust cross-modal alignment remains an open problem in practice^36, 37^. In clinical workflows, neuroradiologists may need to manually cross-reference sequences when robust co-registration is unavailable, which can contribute to cognitive load and interpretive error^38^. Thus, there is significant motivation for an automated, high-accuracy, multi-modal registration approach.

Overall, there is a need for a fast, robust, and accessible MRI preprocessing pipeline that bridges the research-clinic gap. MICAFlow was conceived to fulfill these requirements. MICAFlow is a fully automated MRI preprocessing pipeline that emphasizes accessibility (open-source Pythonic implementation), speed (leveraging machine learning and parallel computing), and clinical relevance of outputs. Unlike research pipelines that prioritize exhaustive analyses at the expense of time, MICAFlow focuses on essential volumetric and dMRI outputs. It incorporates multiple state-of-the-art components: for example, a deep learning segmentation tool, SynthSeg^39^, to drive a novel cross-modal registration method we term Label-Augmented Modality-Agnostic Registration (LAMAReg), and machine learning-based distortion correction for diffusion images. By combining these techniques, MICAFlow achieves robust multi-modal alignment and artifact correction that surpass comparable methods in accuracy. The pipeline is built with modular, reproducible workflow principles (using the Snakemake engine and following the Brain Imaging Data Structure Application (BIDSApp) standard^40^) to facilitate integration into clinical image processing pipelines. In this paper, we describe MICAFlow’s design and innovations in detail, and we report how it compares to existing solutions. We evaluate key performance metrics: processing runtime, registration accuracy across modalities, and data reliability. Results show that MICAFlow can preprocess a multimodal MRI exam in minutes, is comparable to current methods in simple registration problems, and demonstrates significantly improved cross-modal registration fidelity relative to standard algorithms. These results hold in cases with heavy artifacts or distortion and produce outputs with high test-retest reliability across multiple datasets, including in a cohort of patients with epilepsy. Together, these advancements position MICAFlow as a practical bridge between research neuroimaging and clinical practice, enabling clinicians to obtain research-grade images within clinical timeframes.

## Methods

### Pipeline Overview and Architecture

MICAFlow is a comprehensive MRI preprocessing pipeline for T1w, FLAIR/T2w, and dMRI. The pipeline is implemented in Python and orchestrated by the Snakemake workflow engine for modularity, reproducibility, and efficient parallel execution^41^. Using Snakemake, MICAFlow defines all processing steps in a directed acyclic graph structure, which allows automatic parallelization of independent tasks and easy checkpointing/resumption of processing if interrupted. If a long process is halted, MICAFlow can resume from the last successful step. The pipeline adheres to BIDSApp standards for organizing the command line interface and outputs^40^, ensuring compatibility with other neuroimaging tools. Core to the MICAFlow pipeline is LAMAReg, which enables robust multi-modal registrations. A high-level outline of the LAMAReg pipeline is provided with **Figure 1a**, while **Figure 1b** provides an overview of the MICAFlow workflow.

**Figure 1.**
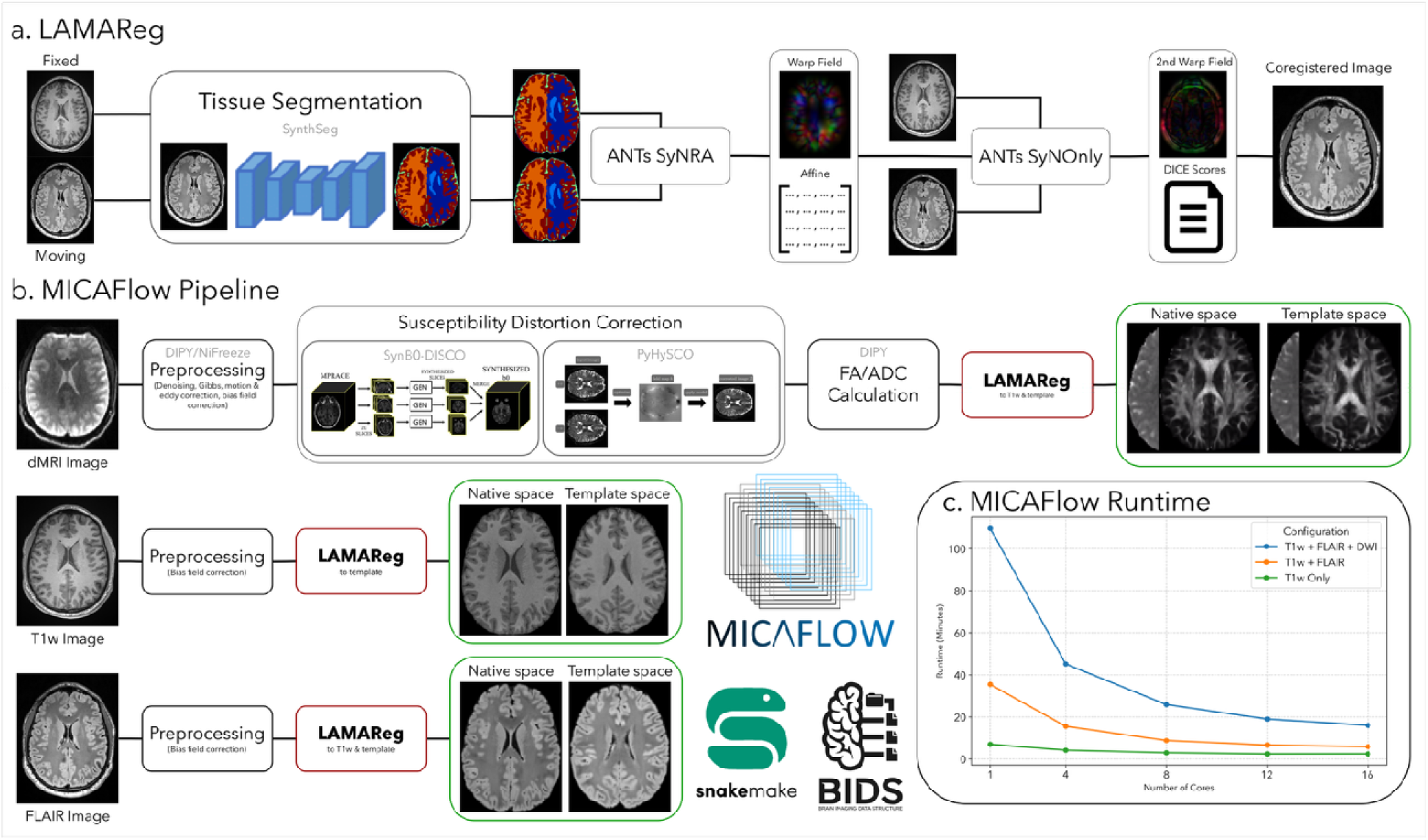
| MICAFlow Workflow and Runtime Performance. **(a)** Diagram of the LAMAReg label-driven registration workflow, illustrated for FLAIR-to-T1w alignment. SynthSeg is used to create label maps from each image, which are then registered via two ANTs SyN stages: a SyNRA (rigid + affine + nonlinear) for initial alignment, followed by a SyN-only deformable refinement on the original intensity images. **(b)** Schematic of the full MICAFlow pipeline. The structural workflow (T1w→MNI and FLAIR→T1w) and the diffusion workflow (dMRI preprocessing and mapping to T1/MNI) are depicted. LAMAReg steps are highlighted in red, and key final outputs (aligned FLAIR, FA/ADC maps in native and MNI152 space) are highlighted in green. Machine-learning based modules for diffusion (Synb0-DISCO/PyHySCO) are noted. **(c)** Pipeline wall-clock runtime as a function of available CPU cores (1, 4, 8, 12, 16 cores) for three input scenarios: full T1w+FLAIR+dMRI (*blue*), T1w+FLAIR only (*orange*), and T1w-only (*green*).

In MICAFlow, the structural MRI branch processes the T1w image and (if available) a FLAIR or T2w image, while the dMRI branch processes the raw dMRI data. Major steps in the structural pipeline include bias field correction, brain extraction, tissue segmentation, cross-modal registration (FLAIR/T2w to T1w), and normalization to stereotaxic space. The diffusion pipeline includes denoising, motion and eddy current correction, EPI distortion correction, diffusion tensor fitting, and registration of diffusion-derived maps to the structural anatomy. Key outputs (highlighted in green in **Figure 1b**) include the bias-corrected T1w image in native and standard (MNI152) space, the aligned FLAIR/T2w in T1w and MNI152 space, and FA/ADC available in native dMRI space, T1w space, and MNI152 space. Throughout the workflow, we leverage multi-core processing: many algorithms such as ANTs^42, 43^ registrations, DIPY diffusion fitting^44^, NiFreeze^45^, PyHySCO^46^, and Synb0-DISCO^47^ are configured to use multi-core processing, and Snakemake runs separate tasks concurrently when possible. This highly parallelized design sigmificantly reduces overall processing time on modern multi-core CPUs.

### SynthSeg-Driven Segmentation and Labeling

A cornerstone of MICAFlow’s design is the use of SynthSeg for robust, contrast-agnostic brain segmentation. SynthSeg is a deep convolutional neural network trained on thousands of artificially generated MRI volumes with varied contrasts and artifacts, enabling it to segment brain tissues reliably across a wide range of imaging sequences^39^. In MICAFlow, we run SynthSeg on each input volume (T1w, FLAIR/T2w, and the b=0 dMRI image) to obtain a labeled brain mask partitioning the image into tissue types (gray matter, white matter, cerebrospinal fluid, ventricles, and various subcortical structures). Importantly, these segmentation maps provide a common anatomical reference that is largely independent of MRI contrast. These label maps serve as stable anchors for downstream processing, particularly for guiding registration, as described next. Using SynthSeg for skull stripping also improves robustness: rather than relying on sequence-specific brain extraction tools, which may be less reliable on atypical contrasts and some pathologies^48^, the pipeline derives brain masks directly from SynthSeg’s segmentation, ensuring consistent brain tissue definition across modalities.

### LAMAReg: Label-Augmented Modality-Agnostic Registration

To achieve reliable alignment between images of different contrasts, we developed LAMAReg, an AI-assisted registration which preserves interpretability by relying on typical analytic registration methods. LAMAReg leverages the SynthSeg segmentations of the moving and target images to drive the registration, instead of relying solely on image intensities. The process is a two-stage deformable registration implemented with the ANTs’ SyN algorithm^43^:

> Stage 1 (Segmentation-to-Segmentation): We first register the label map of the moving image to the label map of the fixed image. This uses ANTs SyN with a full affine + nonlinear transformation (the “SyNRA” configuration: rigid + affine + deformable SyN). Because the inputs are discrete label images representing anatomical structures, the optimizer effectively aligns common regions. This yields an initial transform that brings the two brains into coarse alignment based on their anatomical labels, largely overcoming differences in intensity or artifacts in the original scans.
>
> Stage 2 (Refinement): Next, we fine-tune the alignment by applying a deformable registration on the original images, initialized by the Stage 1 result. In this stage, we use ANTs SyN in deformable-only mode, which refines local alignment while preserving the global correspondence established by the labels. Essentially, this step lets the actual image intensities contribute to the final registration, adjusting for any residual misalignments once the images are roughly in the same space. Due to the registration starting from a strong initialization (Stage 1), the intensity-based optimization generally avoids falling into incorrect local minima that normally plague multi-modal registration^49,50.^

### Structural MRI Preprocessing

For structural images (T1w and FLAIR/T2w), MICAFlow performs a series of conventional preprocessing steps enhanced by the techniques above. First, each image undergoes ANTs’ N4 bias field correction to remove low-frequency intensity inhomogeneities^51^. The T1w image, generally being the highest-resolution anatomical reference, is designated as the primary structural image. After bias correction, the T1w is used as a reference for the LAMAReg-driven FLAIR/T2w and dMRI alignment.

Meanwhile, the T1w image is normalized to MNI152 standard space using LAMAReg for nonlinear template registration, or ANTs for linear template registration. When using LAMAReg for nonlinear template registration, we only use the first stage of the registration, aligning the overall anatomy without directly matching voxel intensities. This design is particularly helpful for template normalization because it prioritizes robust global anatomical correspondence while avoiding potential bias from intensity differences that may be unrelated to spatial alignment. The resulting transform (T1w→MNI) is saved, as it will later be applied to other derivative maps (FA, ADC, FLAIR) so that all outputs can be provided in standardized space.

### Diffusion MRI Preprocessing

MICAFlow’s diffusion preprocessing module handles all steps needed to convert raw dMRI scans into artifact-corrected, anatomically aligned diffusion metrics. The pipeline ingests dMRI volumes with corresponding b-values and gradient directions. We require at least one non-diffusion-weighted volume (b=0) in the series.

### Denoising and Gibbs Correction

As an initial step, we apply basic dMRI preprocessing using DIPY, an open-source diffusion MRI library^44^. This includes denoising with DIPY’s patch2self^52^ algorithm and correction of Gibbs ringing artifacts. These operations improve the quality of dMRI data by reducing high-frequency noise and removing EPI truncation artifacts.

### Motion and Eddy Current Correction

The dMRI series is then fed into NiFreeze, a tool for motion and eddy current correction from the NiPreps group^45^. It removes subject motion by registering all diffusion volumes to a common target and simultaneously accounts for eddy current-induced distortions (which manifest as shear/stretch in diffusion volumes due to strong diffusion gradients). NiFreeze yields a corrected dMRI where all volumes are in geometrical alignment with each other. We take the linear transformation matrices generated during this step and apply them to the gradient direction vectors to ensure they are correctly reoriented relative to the subject’s corrected anatomy. At this stage, the dMRI data is internally consistent but still may be spatially distorted due to susceptibility artifacts.

### Susceptibility Distortion Correction

EPI dMRI is prone to susceptibility-induced geometric distortions along the phase-encoding axis. Correcting these distortions is technically challenging. Common correction strategies often require supplementary acquisitions such as field maps or reverse phase-encoded images, and the distortion field can interact with subject motion^53^. MICAFlow addresses this with a flexible two-pronged approach depending on data availability:

> **Scenario A:** Reverse Phase-Encoding Available - If the clinical dMRI protocol includes a pair of b=0 images with opposite phase-encoding directions (*e.g.,* blip-up/blip-down), we invoke PyHySCO (PyTorch-based Hyperelastic Susceptibility Correction)^46^. PyHySCO uses the two images to estimate the field inhomogeneity-induced displacement field by solving a physics-based model with hyperelastic regularization. It runs on CPU or GPU and can compute the correction in seconds. We apply PyHySCO to compute a deformation that unwarps the diffusion images, yielding a geometrically corrected dMRI volume.
>
> **Scenario B:** No Fieldmap Available - In some clinical scenarios, no reverse-encoded images are acquired due to time constraints. In this case, MICAFlow employs Synb0-DISCO, a deep learning method that synthesizes an undistorted b=0 image from the T1w scan^47^. Essentially, Synb0-DISCO uses the structural image to predict how the b=0 would look if it had no distortion, then MICAFlow calculates the warp needed to make the real b=0 match that synthetic target. While this approach relies on learned priors, it extends distortion correction capability to datasets where traditional corrections are impossible, vastly increasing the clinical utility for standard protocols.

After these steps, the diffusion volumes have been motion-corrected and distortion-corrected. Then, using LAMAReg, we register the corrected b=0 image, now representative of the dMRI series, to the T1w image. This cross-modal registration again uses the label-to-label alignment (b=0 labels to T1w labels) followed by fine intensity alignment. The resulting transform maps the diffusion native space to the T1w native space. We apply this transform to all dMRI-derived maps, ensuring that the diffusion data are aligned with the anatomical image.

### Diffusion Metric Mapping

Once corrections and alignment are complete, we calculate standard dMRI-derived quantitative maps. We fit a diffusion tensor model using DIPY’s reconstruction algorithms to the corrected diffusion data^44^. From the fitted tensor, we generate FA and ADC maps in the native diffusion space. We then apply the previously obtained transforms to bring these maps into the desired spaces: using the LAMAReg transform from dMRI-to-T1w, we resample the FA and ADC maps into T1w native space. This ensures that, for example, an area of high FA can be directly located on the anatomical scan. By concatenating the diffusion-to-T1w transform with the T1w-to-MNI transform, we also warp the FA and ADC maps into MNI152 standard space. This allows group analyses or atlas-based interpretations.

### Healthy Datasets

We evaluated performance of MICAFlow on data from four different healthy MRI datasets. We used a version of the Microstructure Informed Connectomics (MICs) 3T MRI dataset^54^ with an expanded cohort, totaling 144 sessions, each with 40 direction b=700 diffusion shells and reverse-phase encoded images. We also used an expanded cohort of the Precision Neuroimaging and Connectomics (PNI) dataset^55^ comprising 63 sessions at 7T, with b=700 diffusion shells and reverse phase encoding scans. In addition, we used the Epilepsy in Cognition (EpiC) 3T MRI dataset^56^ with 32 sessions, which included b=2000 shells reverse-phase encoding scans. Finally, we used the University of Amsterdam’s test-retest dataset (AmTrT)^57^, which included 68 sessions with b=1000 shells and no reverse-phase encoded images acquired at 3T. For the registration assessments, all 307 sessions were used. For the dataset-wide identifiability analyses, we restricted these sessions to include only the first and second sessions for each subject. A full breakdown of imaging parameters is available in **Table 1**.

**Table 1:**
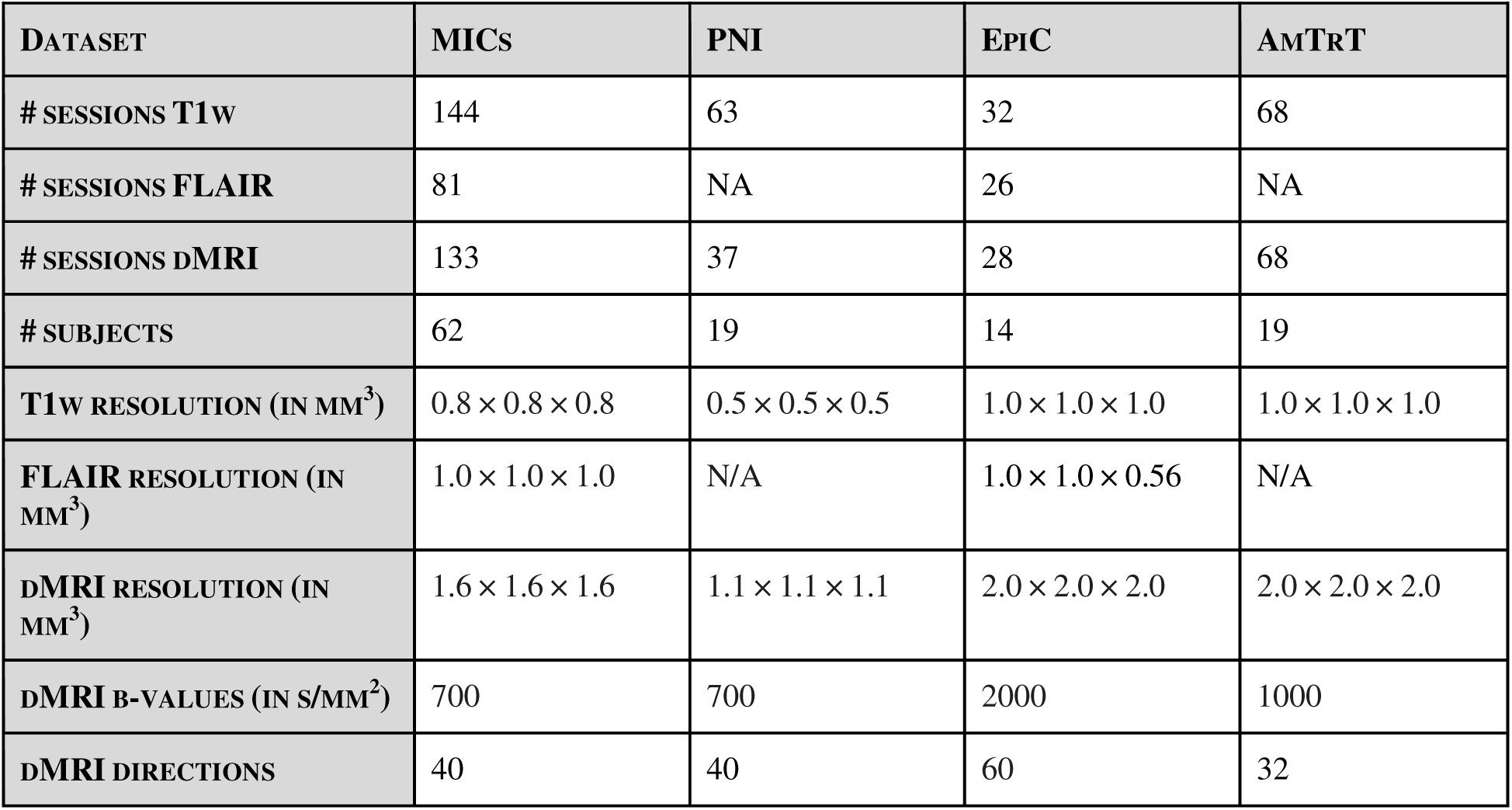
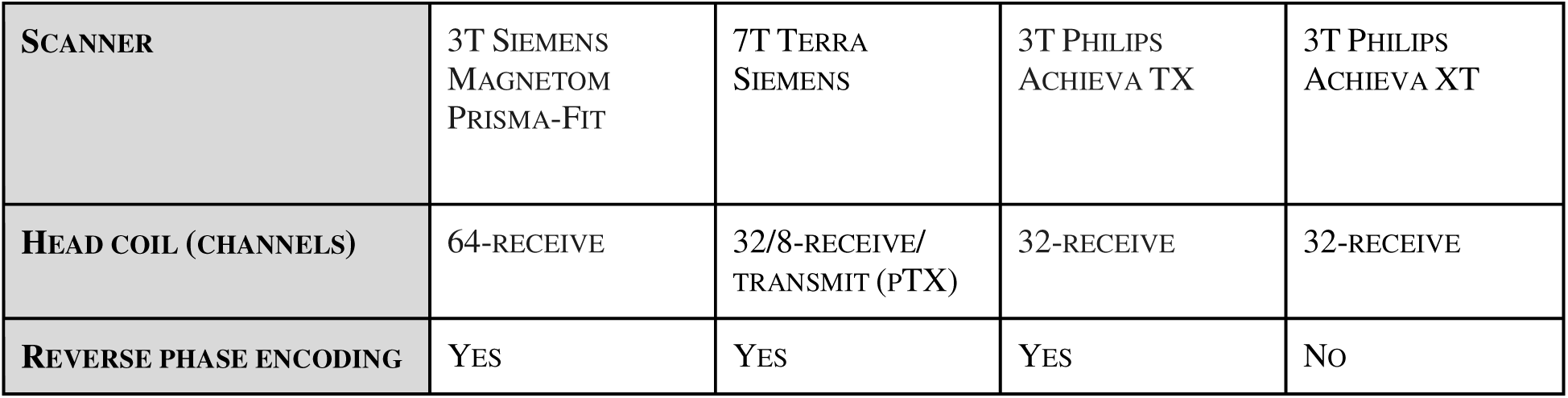

| DATASET | MICs | PNI | EpiC | AmTrT |
| --- | --- | --- | --- | --- |
| # SESSIONS T1w | 144 | 63 | 32 | 68 |
| # SESSIONS FLAIR | 81 | NA | 26 | NA |
| # SESSIONS dMRI | 133 | 37 | 28 | 68 |
| # SUBJECTS | 62 | 19 | 14 | 19 |
| T1w RESOLUTION (IN MM<sup>3</sup>) | $0.8 \times 0.8 \times 0.8$ | $0.5 \times 0.5 \times 0.5$ | $1.0 \times 1.0 \times 1.0$ | $1.0 \times 1.0 \times 1.0$ |
| FLAIR RESOLUTION (IN MM<sup>3</sup>) | $1.0 \times 1.0 \times 1.0$ | N/A | $1.0 \times 1.0 \times 0.56$ | N/A |
| dMRI RESOLUTION (IN MM<sup>3</sup>) | $1.6 \times 1.6 \times 1.6$ | $1.1 \times 1.1 \times 1.1$ | $2.0 \times 2.0 \times 2.0$ | $2.0 \times 2.0 \times 2.0$ |
| dMRI B-VALUES (IN S/MM<sup>2</sup>) | 700 | 700 | 2000 | 1000 |
| dMRI DIRECTIONS | 40 | 40 | 60 | 32 |
| **SCANNER** | 3T SIEMENS<br>MAGNETOM<br>PRISMA-FIT | 7T TERRA<br>SIEMENS | 3T PHILIPS<br>ACHIEVA TX | 3T PHILIPS<br>ACHIEVA XT |
| **HEAD COIL (CHANNELS)** | 64-RECEIVE | 32/8-RECEIVE/<br>TRANSMIT (PTX) | 32-RECEIVE | 32-RECEIVE |
| **REVERSE PHASE ENCODING** | YES | YES | YES | No |

### Clinical Datasets

We evaluated MICAFlow on two datasets comprised of patients with highly varied epilepsy pathologies. The first was a subset of MICs consisting of 60 patients, for whom the most recent two sessions were used. This cohort was balanced by sex (30 female, 30 male), with a mean age of 32.9 ± 12.0 years. Most patients had definite epilepsy by ILAE criteria (58/60), with one likely case and one unclear diagnosis. Epilepsy classification was predominantly focal (55/60), with two generalized cases and three unclear cases. Epileptogenic lateralization was right-sided in 26/60 patients, left-sided in 24/60, bilateral in 4/60, generalized in 2/60, unclear in 3/60, and unavailable in 1/60. The cohort was clinically heterogeneous, although temporal lobe epilepsy was most common, with TLE recorded in 36/60 patients, followed by frontal lobe epilepsy in 8/60, central lobe epilepsy in 2/60, idiopathic generalized epilepsy in 2/60, and smaller numbers of posterior quadrant, neocortical temporal, insular, and multifocal cases. Histopathology was available in a subset of surgical patients and included focal cortical dysplasia, hippocampal sclerosis, gliosis or chronic seizure-related changes, and cavernoma-related pathology. The EpiC patient subset consisted of 19 individuals with temporal lobe epilepsy (15 female; age 31.1 ± 10.6 years). Lateralization was predominantly left-sided (11/19), followed by right-sided (6/19), bilateral (1/19), and unknown (1/19), with mesial temporal sclerosis present in 8/19 patients. The acquisition parameters for both of these datasets are the same as their healthy cohorts, seen in Table 1. All sessions were screened for visible resection cavities and excluded if they were found.

### Evaluation Metrics and Validation Strategy

We evaluated MICAFlow’s processing speed, as well as the quality of the underlying registration. For the evaluation of registration accuracy and robustness we focused on two challenging cross-modal alignments in our dataset, dMRI b=0 to T1w and FLAIR to T1w, as well as a same-modality case (T1w to T1w intra-subject):

#### Runtime Performance

To assess processing speed and scalability, we ran MICAFlow with different input configurations and computational resources. Specifically, we tested three common clinical scenarios: (1) a full multimodal dataset (T1w + FLAIR + dMRI), (2) structural-only (T1w + FLAIR, no diffusion), and (3) T1w only. Each scenario was executed with varying degrees of parallelization: 1, 4, 8, 12, and 16 CPU cores. All runs were on the same commercial hardware (see **Supplementary Table 1** for details). We recorded the total wall-clock time for each run. This experiment mimics how the pipeline would perform in sites with different hardware, from a single-core laptop up to a typical multi-core workstation. **Figure 1c** plots the runtime results as a function of core count for each scenario.

#### Registration Accuracy

The accuracy of image registration was quantitatively evaluated by comparing MICAFlow’s LAMAReg against three widely used registration approaches: *(a)* ANTs SyN with identical parameters to LAMAReg (intensity-based using mutual information)^43^, *(b)* FSL’s epi_reg (a boundary-based registration method designed for dMRI-to-T1w alignment)^13^, and *(c)* FreeSurfer’s EasyReg (a machine-learning powered registration)^58^. Since no ground-truth alignment exists for *in vivo* data, we used similarity metrics as proxies for accuracy. We computed three complementary metrics after each alignment: Mutual Information (MI), an information-theoretic similarity measure that quantifies the statistical dependence, or shared information, between image intensities in the two images^59^; we use the negative MI, so lower values indicate better alignment. Modality Independent Neighbourhood Descriptor (MIND) dissimilarity, a local structural measure derived from MIND^60^ which is based on local self-similarity and is designed to capture neighbourhood structure that is preserved across modalities; when the distance between MIND descriptors is reported, lower values indicate closer local structural correspondence. Normalized Gradient Fields (NGF) is a multimodal registration measure that emphasizes agreement between image edges and contour directions; when formulated as an NGF distance or error term, lower values indicate better boundary alignment^61^. These metrics capture different aspects of registration quality: MI is global and tied to overlapping intensity distributions, MIND evaluates local texture alignment, and NGF emphasizes edge alignment. For each registration method and image pair, we computed these metrics after alignment, and performed paired t-tests followed by FDR correction^62^.

#### Robustness to Artifacts

To assess the robustness of each registration approach to common scanning artifacts, we introduced synthetic noise and motion artifacts into the images using TorchIO^63^ and evaluated registration performance under these degraded conditions. In each case, the fixed T1w image was left alone, and the moving image (T1w, FLAIR, or b=0) was perturbed with simulated artifacts before registration. Noise was added by injecting Gaussian noise at different levels, sampling from a distribution with variance as a percentage of the 95% intensity range of the base image. Motion artifacts were simulated by applying an in-plane rotation and translation to the moving image^63^. We then applied each registration method to align the artifact-degraded image to the reference T1w. After registration, we computed the same quality metrics (MI, MIND, NGF) between the T1w and undegraded image, warped by the transforms calculated on the degraded image, to quantify alignment accuracy under each artifact condition. This experiment was repeated for all eligible subjects across the four datasets, providing a distribution of metric outcomes for each method and condition. By comparing these results, we could evaluate how robust each method’s performance was when images were noisier or initially misaligned.

#### Data Reliability (Identifiability)

To determine whether MICAFlow’s preprocessing yields more reliable and subject-consistent data, we adopted the framework of brain “fingerprinting” or identifiability. The concept of identifiability is that a good preprocessing pipeline should maximize the similarity between two scans of the same subject while minimizing the similarity between scans of different subjects^64^. We used test-retest datasets (each containing subjects scanned on two or more separate sessions) processed through MICAFlow. For each dataset, we computed the correlation between every paired set of multimodal scans across subjects for a given modality or feature. When combining multiple modalities for correlation, before all masked data was collapsed to a vector, z-scoring was done on each image. We then calculated the identifiability, defined as the difference between the mean within-subject correlation and the mean between-subject correlation multiplied by 100, as previously described^64^. A higher identifiability score indicates that repeat scans from the same individual are much more alike than scans from different individuals, meaning the pipeline preserved each person’s unique brain signature while filtering out random noise. In our analysis we compressed all modalities into a single feature vector after brain masking, computing identifiability for each dataset under two preprocessing conditions: linear (affine-only) template registration vs. nonlinear (SyN) template registration. Additionally, we computed the registration quality metrics (MI, MIND, NGF) for key alignments in each dataset to compare the alignment fidelity across datasets and registration approaches. To assess the reproducibility of MICAFlow’s dMRI processing, we calculated the intra-subject correlation of DTI maps, using a once-eroded white matter mask, calculated with SynthSeg in nonlinear MNI152 space. All statistical analyses were performed in Python, and significance levels are indicated in the figures.

#### Clinical Utility

In order to assess the ability for MICAFlow to operate on cases with pathology, we leveraged cases with epilepsy matched to the exact same acquisition parameters as our healthy cohorts. We ran MICAFlow on these cases, and compared identifiability of patients and controls, after running both linear and nonlinear template registration. While there is no ground truth for the successful template registration of pathology, we measure this by proxy using the intra-subject correlation of all scans. We extracted all voxel values within a brain mask from the most recent two sessions of each subject which included T1w, FLAIR, and dMRI, and performed a Pearson correlation on the resulting vectors. When combining multiple modalities for correlation, before all masked data was collapsed to a vector, z-scoring was done on each image. This was performed on the linear and nonlinear template registration, for patients and controls, in both the MICs and the EpiC datasets.

### Code Availability Statement

MICAFlow is available as an open-source Python package. The full source code, version history, and issue tracker are hosted on GitHub https://github.com/MICA-MNI/MICAFlow, with documentation maintained on ReadTheDocs https://mica-mni.github.io/MICAFlow. Command-line usage and options are documented via the built-in help interface (*i.e.*, micaflow --help). The package is also distributed via PyPI https://pypi.org/project/MICAFlow/ and can be installed with “pip install micaflow” on systems with a supported Python environment (≥3.10). Models for Synb0-DISCO and MNI152 atlases are downloaded at runtime (repository link: https://github.com/MICA-MNI/MICAFlow-Models).

LAMAReg is additionally released as a standalone open-source package, with source code hosted on GitHub https://github.com/MICA-MNI/LAMAReg and distribution via PyPI https://pypi.org/project/LAMAReg. Command-line usage and options are also documented via the built-in help interface (*e.g.,* lamareg --help). LAMAReg uses the original SynthSeg model weights; for lightweight installation, the weights are hosted as a set of packaged components that are downloaded and reassembled at runtime (https://github.com/MICA-MNI/LAMAR-Models).

## Results

On CPU-only hardware, MICAFlow scaled efficiently with available cores (**Figure 1c**): for a full multimodal session (T1w+FLAIR+dMRI), runtime decreased from ∼110 min (1 core) to ∼17 min (16 cores). Structural-only configurations were faster (∼6 min for T1w+FLAIR; ∼3 min for T1w-only on 16 cores). On 4 cores, the multimodal pipeline ran in <45 min and the structural pipeline in ∼10 min, and on typical 12-16-core workstations a comprehensive exam finished in ∼12-18 min without requiring GPU acceleration. Optional GPU availability can further accelerate specific modules but it is not necessary to achieve the reported end-to-end runtimes. On all the cases tested in this paper, no manual intervention was needed for any pipeline stage; we observed no MICAFlow processing failures.

Using paired *t*-tests with Benjamini-Hochberg FDR correction, LAMAReg generally produced lower registration error than the conventional baselines across MI, MIND, and NGF. In the within-modality T1w→T1w condition, LAMAReg outperformed EasyReg on MI, MIND, and NGF (all *p_FDR_*<0.001), and outperformed ANTs on MIND (*p_FDR_*=0.003), but did not differ significantly from ANTs on MI (*p_FDR_*=0.289) or NGF (*p_FDR_*=0.062). In the FLAIR→T1w task, LAMAReg outperformed EasyReg on MI (*p_FDR_*<0.001) but did not differ from ANTs on MI (*p_FDR_*=0.560). For MIND, LAMAReg outperformed ANTs (*pFDR*<0.001) but was not different from EasyReg (*p_FDR_*=0.719). For NGF, LAMAReg outperformed both ANTs (*p_FDR_*<0.001) and EasyReg (*p_FDR_*=0.031). In the dMRI (b=0)→T1w task, LAMAReg outperformed ANTs, FSL, and EasyReg on all three metrics (all *p_FDR_*≤0.001). Full results are available in **Supplementary Table 2**. Overall, LAMAReg provided the strongest and most uniform improvements in the challenging dMRI-T1w setting.

**Figure 3** shows that registration performance degrades with increasing noise and simulated motion artifacts across all methods, but the magnitude and consistency of degradation vary by modality and metric. For T1w→T1w, LAMAReg exceeded EasyReg across almost all metrics, whereas ANTs became competitive under stronger perturbations, including a significant advantage at 10% noise (Δ0.08 MI, Δ0.0012 MIND, Δ0.008 NGF, *p_FDR_*<0.001). EasyReg notably had a significant advantage on the MI metric at 10% noise as well (Δ0.029 MI, *p_FDR_*<0.001). For FLAIR→T1w, LAMAReg consistently either outperformed or was not significantly different from ANTs across artifact levels, while differences versus EasyReg were more variable; at the strongest noise level, ANTs showed slightly lower MI than LAMAReg but without a significant separation (Δ0.012 MI, *p_FDR_*=0.054). EasyReg was found to outperform LAMAReg on the MIND metric at the 2.5% noise level (Δ2.12e^-5^, *p_FDR_*=0.029), as well as at 5 degrees (Δ5.33e^-5^, *p_FDR_*=0.003) and 10 degrees (Δ5.22e^-5^, *p_FDR_*=0.008) of motion. For dMRI (b=0)→T1w, LAMAReg remained the most stable overall, achieving the lowest error for MI, MIND, and NGF across all artifact severities. The full results are available in **Supplementary Table 3**. Overall, the robustness benefit of LAMAReg is most pronounced for challenging cross-contrast alignment, particularly dMRI→T1w.

**Figure 2.**
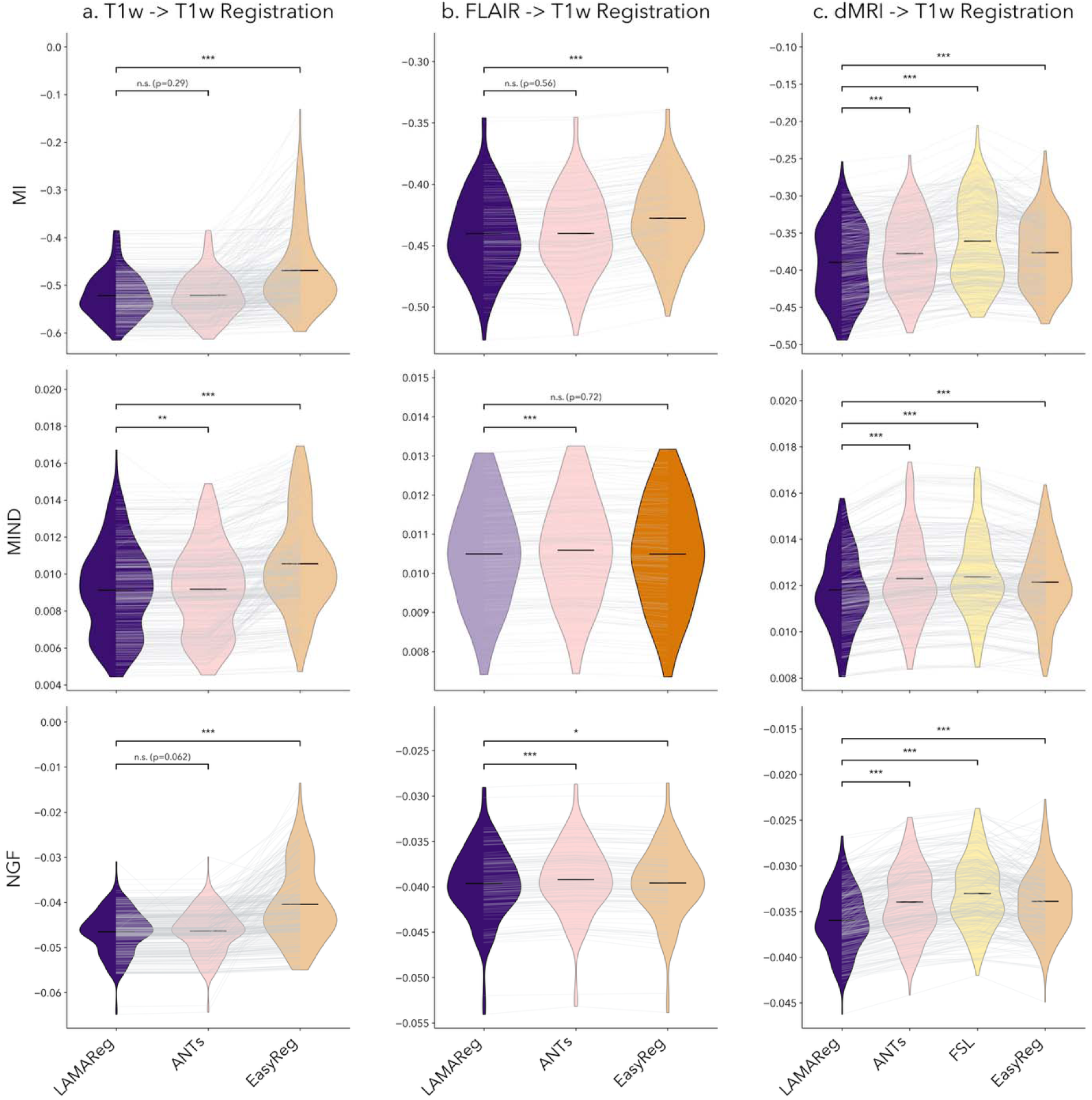
|. Comparative registration accuracy of LAMAReg vs. conventional methods. Violin plots show the distribution of registration quality metrics for different methods across three alignment tasks: **(a)** T1w to T1w (intra-subject alignment between two scans of the same subject), **(b)** FLAIR to T1w, and **(c)** dMRI b=0 to T1w. MICAFlow’s LAMAReg (Purple) is compared to standard ANTs SyN (Pink), FSL’s epi_reg (Yellow), and FreeSurfer’s EasyReg (Orange). Three metrics are reported (lower values or more negative values indicate better alignment): Mutual Information (MI), Modality Independent Neighbourhood Descriptor (MIND), and Normalize Gradient Fields (NGF). The method with the lowest (best) average error for each metric is highlighted, while the others are more transparent. Asterisks denote statistical significance of LAMAReg’s improvement over others (*=*p*<0.05; **=*p*<0.01; ***=*p*<0.001, corrected for multiple comparisons).

**Figure 3.**
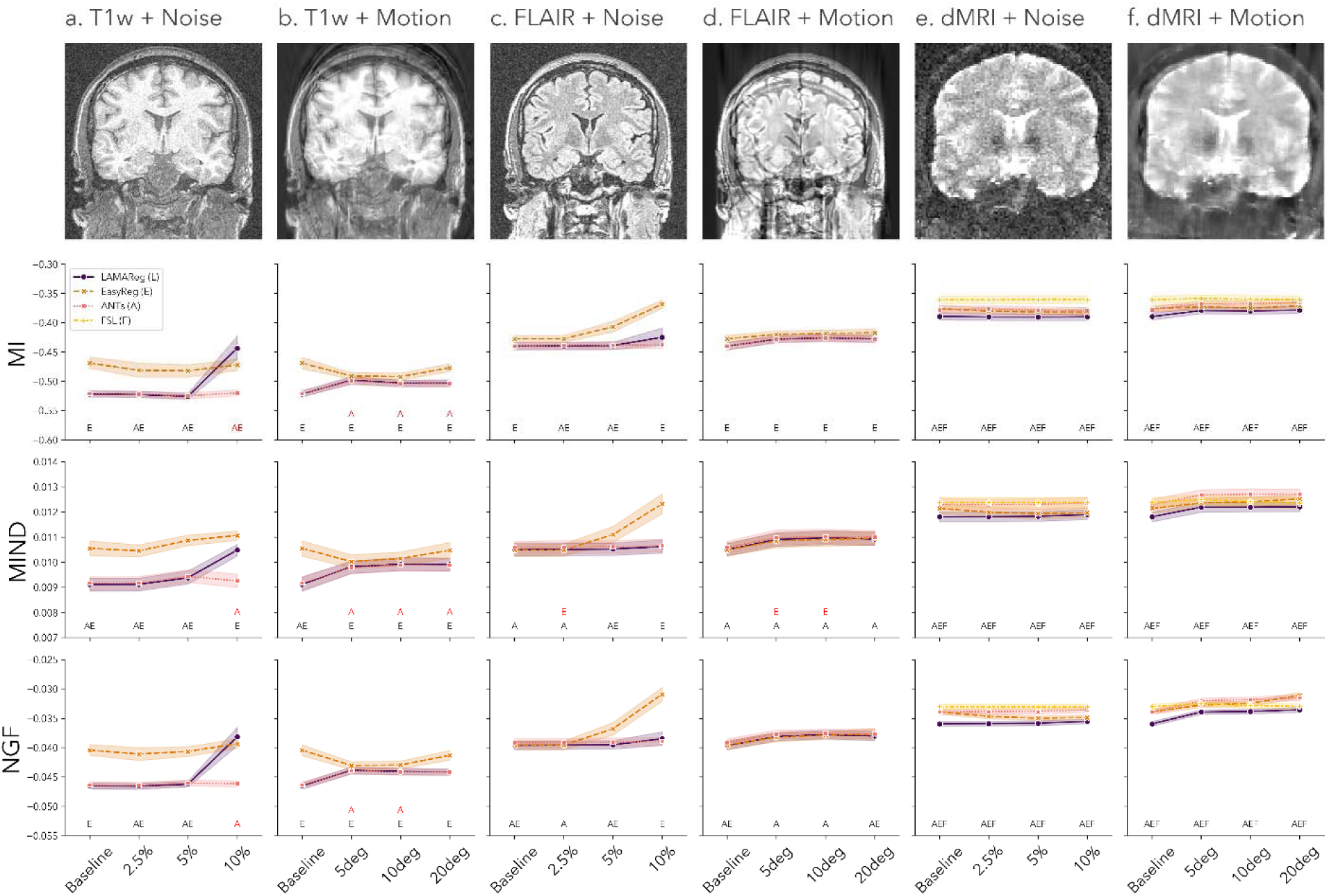
|. Robustness of registration to noise and motion. Representative T1w, FLAIR, and dMRI (b=0) images are shown at baseline and after additive Gaussian noise (2.5-10%) or simulated in-plane motion (5-20° rotation an translation). Registration quality (MI, MIND, NGF; lower is better) is reported at each artifact level across subjects/sessions, with pairwise comparisons evaluated using paired t-tests with Benjamini-Hochberg FDR correction. Data here is aggregated across all datasets. Columns **(a, b)** depict results for T1w→T1w registration wit added noise and motion artifacts. The same is shown for FLAIR→T1w in columns **(c, d)**, and for dMRI (b=0)→T1w in columns **(e, f)**. One-letter abbreviations for each method are shown at the bottom of each figure. Letters in black indicate the methods which LAMAReg was significantly outperforming, while letters in red show which methods significantly outperformed LAMAReg.

**Figure 4.**
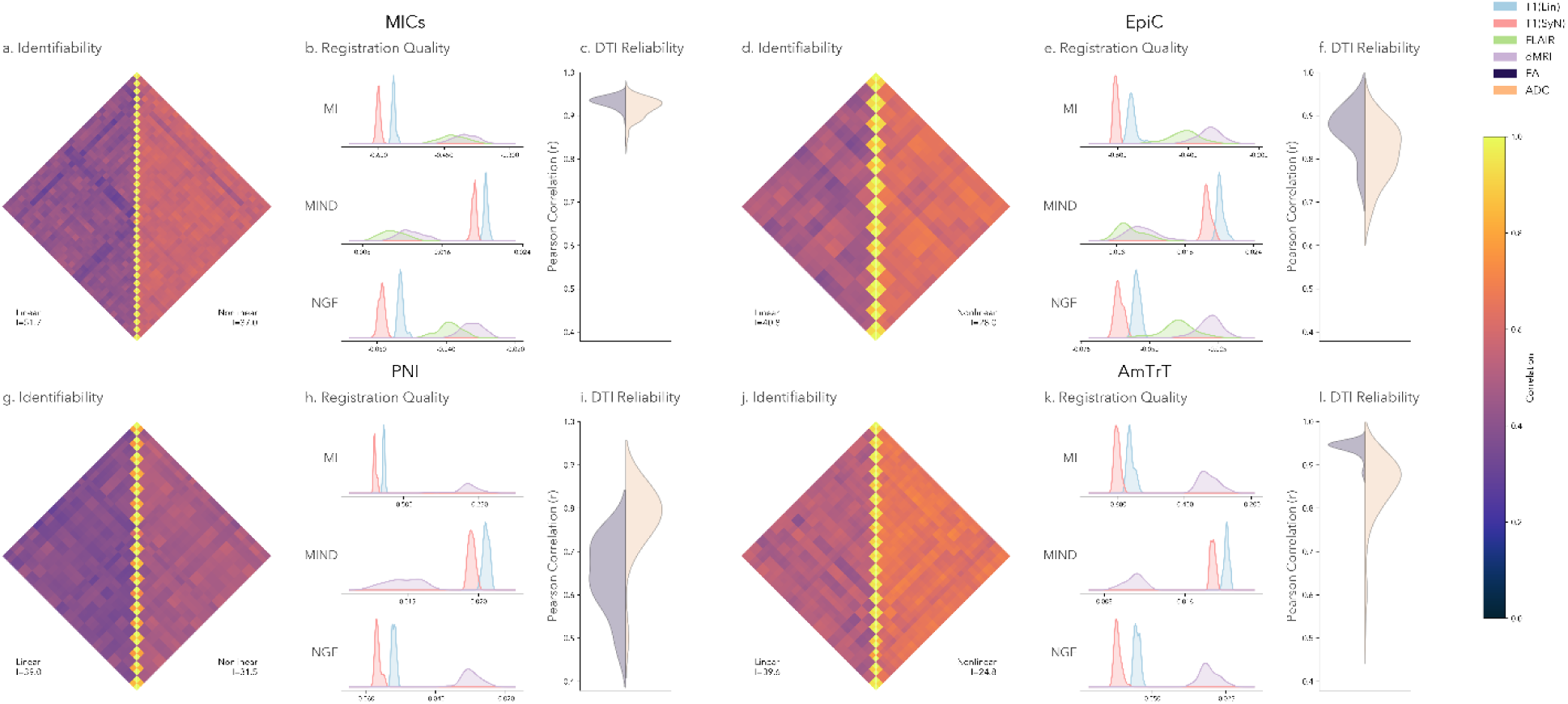
|. Consistency of MICAFlow outputs across different datasets and registration approaches. **(a)** Correlation “fingerprint” matrices for an example dataset (MICs), showing Pearson correlation between every pair of subjects’ multimodal images after linear affine-only registration (*left*) and after full nonlinear (SyN) registration (*right*) t MNI space. Each matrix is ordered by subject. **(b)** Distributions of registration quality metrics (MI, MIND, NGF) for key registration tasks under MICAFlow, aggregated across four independent datasets (MICs, EpiC, PNI, AmTrT). Blue and red correspond to T1w→MNI alignment using linear vs nonlinear (SyN) registration, respectively; green corresponds to FLAIR→T1w alignment, and purple to dMRI b=0→T1w alignment. **(c)** Violin plot showing the intra-subject correlation of dMRI DTI metrics in white matter, FA (purple) and ADC (orange). Panels **(d, e, f)**, **(g, h, i)**, and **(j, k, l)** show the same analyses for the other three datasets.

**Figure 5.**
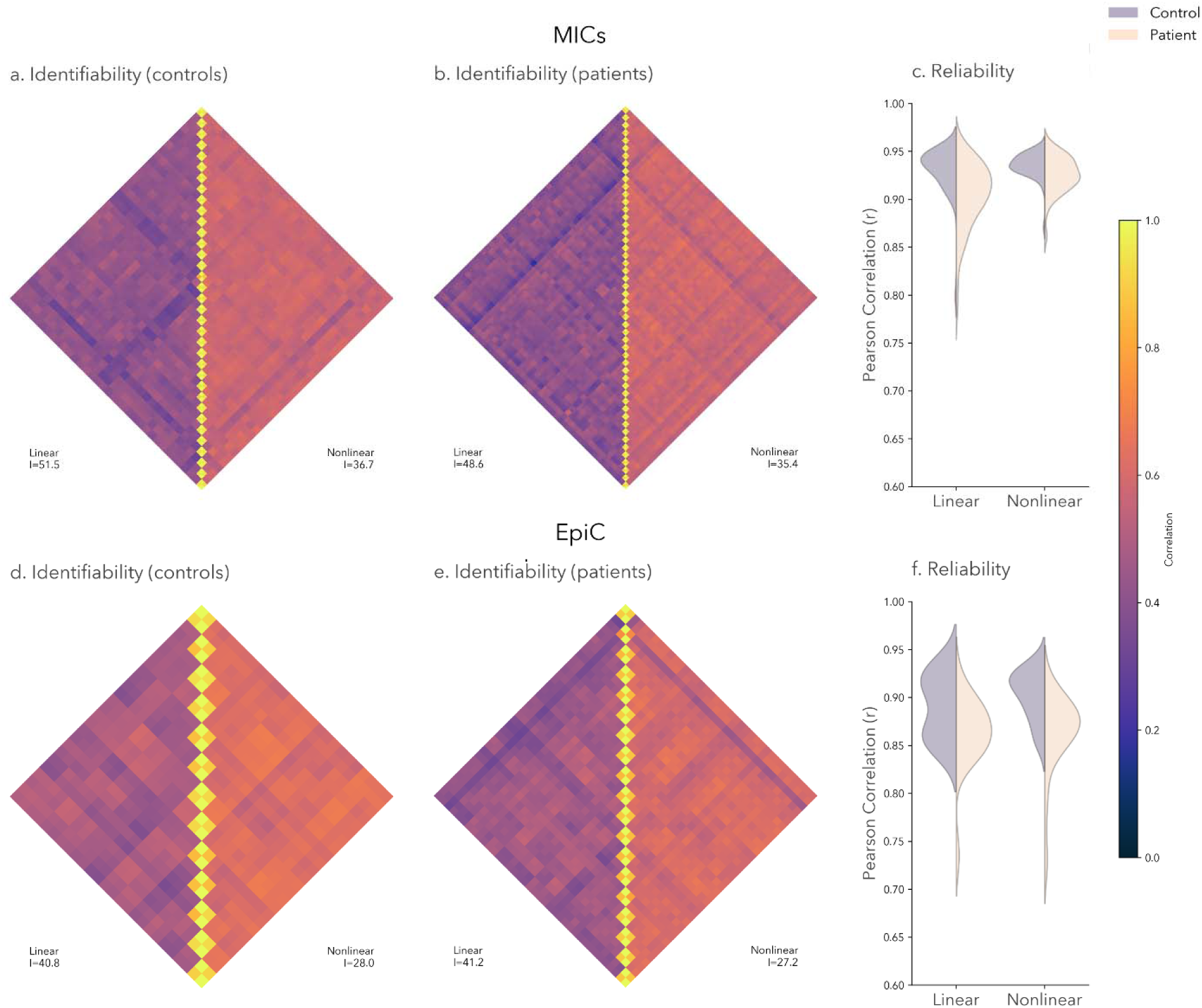
|. Consistency of MICAFlow outputs in clinical populations. (**a)** Correlation matrix for the MICs dataset showing Pearson correlation between every pair of subjects’ multimodal images after linear affine-only normalization (*left*) and after full nonlinear (SyN) normalization (*right*) to MNI space. Each matrix is ordered by subject. **(b)** Correlation matrix for the patient cohort of the MICs dataset, in the same format as the previous **(c)** Intra-subject correlations of both patients (*purple*) and controls (*orange*) after linear (*left*) and nonlinear (*right*) template registration. **(d, e, f)** Same results seen in the EpiC dataset.

Across all four cohorts (MICs, EpiC, PNI, AmTrT), MICAFlow produced strongly subject-identifiable T1w outputs under both affine-only and nonlinear (SyN) normalization, with consistent identifiability. However, identifiability was systematically higher with affine normalization (range: 39-52) than with SyN (25-37), indicating a reproducible trade-off in which nonlinear warping increases anatomical standardization while reducing subject-specific discriminability. This pattern was consistent across datasets, including the example MICs cohort (51.7 linear vs 37.0 SyN). The full results are included in **Supplementary Table 4.**

Registration quality metrics showed high cross-dataset consistency in both absolute values and relative ordering. For T1w→MNI, SyN yielded uniform improvements over affine normalization in every dataset and for all three metrics, with similar means across cohorts (e.g., linear MI - 0.565 to -0.559 vs SyN MI -0.605 to -0.593; linear MIND 2.01e^-2^-2.05e^-2^ vs SyN MIND 1.86e^-2^- 1.95e^-2^; linear NGF -0.055 to -0.052 vs SyN NGF -0.061 to -0.057). The magnitude of SyN’s gain was similarly stable across datasets (ΔSyN-linear≈-0.030 to -0.046 for MI, -1.03e-3 to - 1.52e-3 for MIND, and -0.005 to -0.007 for NGF). Cross-modal registrations exhibited the expected shift toward less favorable MI and NGF compared with T1w→MNI while maintaining low MIND values. Notably, dMRI (b=0)→T1w metrics fell within narrow ranges across all datasets (MI -0.40 to -0.28; MIND 0.011-0.015; NGF -0.032 to -0.028), and FLAIR→T1w (available in MICsand EpiC) was similarly consistent (MI -0.429 to -0.411; MIND 9.5e^-3^-1.10e^-2^; NGF≈-0.039). The full results of dataset-wide registration quality are available in **Supplementary Table 5**. Collectively, the reproducible metric ranges and invariant ordering across cohorts indicate that MICAFlow’s outputs and registration behavior are stable across heterogeneous datasets, though the 7T PNI dataset showed lower DTI reproducibility.

The reproducibility of DTI metrics varied across the datasets, with the MICs dataset having the highest intra-subject correlation of DTI metrics (FA: 0.931, ADC: 0.917). The PNI dataset had relatively low reproducibility (FA: 0.643, ADC: 0.779), likely due to more marked dMRI artifacts seen at 7T. In the EpiC dataset, reproducibility was generally high (FA: 0.867, ADC: 0.813), but still had greater variability than MICs, which may be caused by the greater b-values (2000 versus 700, respectively). The AmTrT dataset had high reproducibility for the FA metrics (0.941), but had notably lower and far more variable reproducibility for ADC (0.849). For all the datasets other than PNI, ADC had a lower reproducibility than FA. The PNI dataset’s low reliability and inconsistency in this pattern may be due to high signal dropout and noise in the original images. The full DTI reproducibility metrics can be seen in **Supplementary Table 6**.

Across both clinical datasets, MICAFlow preserved high intra-subject similarity in patients with epilepsy, despite expected anatomical and pathological variability. Overall, intra-subject correlations were substantially higher than inter-subject correlations (mean r=0.913 ± 0.035 vs. r=0.494 ± 0.089), indicating that repeat scans from the same participant remained strongly identifiable after preprocessing. In MICs, controls showed high reproducibility after both linear (r=0.934 ± 0.019) and nonlinear registration (r=0.936 ± 0.013). Patient scans were modestly lower but remained highly reproducible, with mean correlations of r=0.915 ± 0.030 after linear registration and r=0.925 ± 0.019 after nonlinear registration. In EpiC, correlations were lower overall but remained robust: controls showed mean correlations of r=0.896 ± 0.030 after linear registration and r=0.902 ± 0.025 after nonlinear registration, while patients showed mean correlations of r=0.863 ± 0.039 and r=0.863 ± 0.041, respectively. The full reliability metrics are available in **Supplementary Table 7.**

Consistent with the healthy-cohort analyses, identifiability scores were higher after affine-only normalization than after nonlinear SyN normalization, reflecting the same trade-off between preserving subject-specific imaging signatures and improving anatomical standardization. In MICs, identifiability decreased from 51.5 to 36.7 in controls and from 48.6 to 35.4 in patients after nonlinear registration. In EpiC, the same pattern was observed, with identifiability decreasing from 40.8 to 28.0 in controls and from 41.2 to 27.2 in patients. Together, these results suggest that MICAFlow remains stable in epilepsy cohorts and can preserve subject-specific multimodal MRI structure even in the presence of clinical pathology, while nonlinear normalization provides greater anatomical standardization at the cost of reduced individual discriminability. The full identifiability metrics are available in **Supplementary Table 8.**

## Discussion

MICAFlow is a fully automated, lightweight, and fast MRI preprocessing pipeline designed to translate advanced neuroimaging techniques into clinical and research settings. It processes and aligns structural (T1-weighted, FLAIR/T2w) and diffusion (dMRI) scans within a single workflow. The pipeline uses machine-learning derived segmentation for robust, contrast-agnostic brain registration and performs modality-specific preprocessing steps in parallel. Outputs include skull-stripped T1w images, computed dMRI metrics, and aligned volumes in the native, T1w, and MNI spaces. MICAFlow is designed for accessibility, being able to complete a full multimodal exam in under an hour on a ubiquitous 4-core workstation. Snakemake and multi-threaded tools are used to parallelize tasks across cores. This design makes MICAFlow feasible for time-constrained clinical workflows. MICAFlow introduces LAMAReg, a label-augmented, modality-agnostic registration method. LAMAReg leverages anatomical segmentations as priors to align images across contrasts. In practice, it achieves significantly better alignment of FLAIR and diffusion images to T1w than conventional methods, while remaining robust to common artifacts. Across diverse cohorts, MICAFlow maintains consistent performance. Subject-identifiability tests and registration metrics are stable across datasets, indicating high reproducibility. By combining machine learning and accessibility-focused design, MICAFlow dramatically reduces preprocessing time and complexity. It bridges the gap between research-grade analysis and clinical practice, enabling widespread adoption of dMRI without specialized hardware or long runtimes.

MICAFlow has two parallel branches. The structural branch processes T1w (and optional FLAIR/T2w) images: it performs brain extraction, SynthSeg-driven tissue segmentation, FLAIR/T2w→T1w registration, and spatial normalization to MNI space. The diffusion branch processes raw dMRI data: it applies DIPY for denoising, NiFreeze for motion and eddy-current correction, uses Synb0-DISCO for EPI distortion correction if reverse-phase encoding scans are not available, fits the diffusion tensor, and registers the dMRI volume to the T1w image. All steps run under Snakemake, with multi-threaded tools executing in parallel to maximize efficiency. This modular, parallel design dramatically reduces total processing time on multi-core CPUs. MICAFlow scales efficiently with available CPU cores. For example, a 4-core workstation completes an exam in ≈45 minutes, an 8-core machine in ≈30 minutes, and a 12-core workstation in ≈18 minutes. Even with standard desktop hardware, the full pipeline finishes in under an hour. These results confirm that MICAFlow’s multi-core, commodity-hardware design makes same-day processing feasible without specialized equipment. Contemporary pipelines including dMRI processing such as QSIPrep^14^ and DESIGNER/DESIGNER-v2^65^ are highly precise at the cost of speed. DESIGNER-v2 has been reported to complete comparable preprocessing steps in approximately 52 minutes on a 24-core server with 256 GB of memory. Direct runtime comparison with QSIPrep is difficult because runtime is not consistently reported and depends strongly on acquisition, configuration, and hardware; however, QSIPrep is designed as a more comprehensive diffusion preprocessing framework and may therefore require longer processing times in many settings. MICAFlow is reported here to have a faster runtime than DESIGNER by ≈7 minutes while using only 4 cores on a consumer-grade workstation, though exact comparison is not possible due to a lack of workstation specifications.

We compared MICAFlow’s LAMAReg to standard registration methods ANTs SyN, FreeSurfer’s EasyReg, and FSL’s epi_reg using three metrics. Mutual Information (MI) measures global intensity overlap. MIND generally detects the overlap of intensity patterns in the image, and Normalized Gradient Fields (NGF) measure local structural and edge alignment. Lower values of MIND/NGF generally indicate better alignment of fine structures. We found that LAMAReg outperformed all three methods on the challenging cross-modal case of dMRI-T1w registration^32^. These results position LAMAReg as a competitive option for dMRI-T1w coregistration. For the other cross-modality case, FLAIR-T1w, we showed that LAMAReg beats EasyReg with MI, which is to be expected since LAMAReg is optimizing MI, while it is not significantly different from ANTs. For MIND, we see that LAMAReg significantly outperforms ANTs, while being comparable to EasyReg, and for NGF, we see that LAMAReg significantly outperforms both ANTs and EasyReg. While the relationship here is less strong than with dMRI, we still see that LAMAReg maintains a lead over the other two registration methods. For intra-subject T1w registration, we found that LAMAReg consistently outperforms EasyReg on all metrics, while showing no significant difference compared to ANTs on MI and NGF, and outperforming it on MIND. One interesting finding is that the registration metrics for EasyReg showed more variability, suggesting a high number of misregistrations. By incorporating tissue labels, LAMAReg consistently achieves more accurate multimodal registration than standard approaches. The results are consistent with the idea that tissue-label priors provide stronger anatomical guidance and can reduce failure modes of purely intensity-based multimodal registration, which is known to be susceptible to poor local optima because mutual-information-based matching lacks explicit spatial information^66, 67^.

We evaluated how registration accuracy degrades under simulated imaging artifacts. Random Gaussian noise (0-10% variance) and rigid head motion (0-20 mm translations and 0-20 degree rotation) were added to the input T1w, FLAIR, and dMRI images. We then measured the relative change in alignment metrics (MI, MIND, NGF) compared to the clean (baseline) case. Performance seemed to generally degrade along with the strength of artifacts. LAMAReg performed exceptionally well on the dMRI registration cases, consistently being the top performing method. For FLAIR, LAMAReg showed strong performance, only occasionally having a significantly worse value at low values of noise and motion artifacts on the MIND metric compared to EasyReg. Interestingly, we see a rapid and consistent degradation in the registration quality from EasyReg as a result of noise addition. This is especially interesting because this same effect is not seen in other modalities. Due to EasyReg being a machine-learning method, it suggests that it was not trained on noisy FLAIR images. Our intra-subject T1w registration shows that LAMAReg robustly outperforms other registration algorithms at lower noise levels, but at our highest noise level, we see a spontaneous degradation in registration quality. This represents the risk inherent to segmentation-based registration; where the underlying segmentation fails, the registration will also fail. Fortunately, this also makes registration errors easy to detect. We see a similar issue in the high noise condition on FLAIR registration, although it is far less pronounced. For the intra-subject T1w motion artifacts, we see that using default ANTs SyN registration generally outperformed LAMAReg by a small margin, likely indicating that on T1w scans with motion artifacts, our segmentations were not as accurate. A minor, but surprising finding was that many of the registration methods, but especially EasyReg, performed better when low levels of noise (2.5%) were added. This is known in the case of MI-optimizing registration algorithms^68^ as a way of overcoming the rugged optimization landscape, but has not been reported in machine-learning based algorithms like EasyReg.

We tested MICAFlow on four diverse datasets (MICs, EpiC, PNI, AmTrT) with different scanners and populations. The goal was to assess consistency of outputs and registration across cohorts. Our evaluations highlight a consistent trade-off between anatomical standardization and subject discriminability across all four cohorts: nonlinear template registration improved conventional registration metrics for T1w→MNI alignment in every dataset, but identifiability scores were systematically higher after affine-only normalization (roughly 39-52) than after SyN warping (roughly 25-37), because within-subject similarity remained high in both cases whereas nonlinear normalization also increased between-subject similarity and therefore reduced the separation between each subject’s “self” and “others” correlations. This pattern is highly consistent with the fingerprinting literature, in which subject identifiability is explicitly defined by the gap between within-subject and between-subject similarity^64, 69^, and with prior structural MRI work showing that brain anatomy contains stable individual-specific information that can support subject identification across repeated scans^70–72^. In addition, because the purpose of spatial normalization is to reduce inter-subject anatomical variability in common space^73^, it is plausible that more aggressive nonlinear warping can improve alignment while partially attenuating the individual-specific variation that contributes to identifiability. This interpretation also fits the broader dMRI reproducibility literature, which has generally found that FA- and ADC-derived measures can be stably reproduced across sessions when preprocessing is well controlled^74, 75^. Additionally, we saw that MICAFlow retained generally high intra-subject correlation of dMRI-derived metrics, with a notable exception on the 7T PNI dataset. dMRI at 7T is still an evolving field, with numerous challenges not seen in lower-field dMRI^76^. We also observed a fairly high reproducibility in the AmTrT dataset, notable due to its lack of reverse phase-encoding images, which can pose a challenge to tensor fitting^77, 78^. Taken together, our results suggest that nonlinear registration is preferable when the goal is maximal anatomical correspondence in standard space, whereas affine normalization may better preserve subject-specific imaging fingerprints for longitudinal tracking or individual-level matching.

We next extended these analyses to clinical data from heterogeneous epilepsy cohorts. Because ground-truth registrations are generally unavailable in clinical MRI, we used intra-subject correlation across repeat sessions as a surrogate measure of processing stability. A limitation of this approach is that, in long-standing epilepsy, differences between timepoints may partly reflect disease-related changes in brain structure rather than preprocessing variability alone^79^. Across the MICs and EpiC cohorts, MICAFlow preserved strong intra-subject similarity in patients, with only modest reductions relative to controls, despite heterogeneous epilepsy syndromes and pathologies including mesial temporal sclerosis, focal cortical dysplasia, gliosis, chronic seizure-related change, and cavernoma-related abnormalities. This is a demanding validation setting because clinical MRI data often depart from the assumptions under which image-processing methods are developed: pathology can alter tissue contrast and anatomical correspondence^80^, motion^81^ and acquisition variability^82^ can degrade image quality, and homogeneous development datasets may limit real-world generalization^83^. These issues are particularly relevant in epilepsy, where subtle epileptogenic abnormalities can be difficult to visualize on structural MRI, and where multicentre lesion-detection studies have emphasized the need for robust processing and modelling across heterogeneous scanners, protocols, and pathology types^84, 85^. The preservation of strong patient identifiability after both affine and nonlinear normalization suggests that MICAFlow generalizes beyond healthy or curated research data and can preserve individualized multimodal structure in clinically heterogeneous epilepsy cohorts. However, the lower correlations observed in EpiC and the reduction in identifiability after nonlinear warping indicate that pathology-aware validation remains essential. More aggressive normalization may improve anatomical correspondence, but, as observed here, it can also reduce subject-specific signal and may be less reliable when lesions, abnormal tissue contrast, or imperfect correspondence violate the assumptions of standard registration approaches^80, 86, 87^. Together, these results support MICAFlow as a practical preprocessing framework for clinical neuroimaging studies, where robustness to pathology and acquisition variability is as important as performance on conventional registration benchmarks.

The current pipeline focuses on structural (T1w, FLAIR/T2w) and single-shell dMRI. It does not process other modalities such as functional MRI, perfusion imaging, or advanced multi-shell diffusion at this point. Including such modalities would require additional development and broaden the pipeline beyond the core structural and diffusion workflows emphasized here. What is presented here can serve as a foundation onto which multi-shell and additional modalities can be added. Although functional MRI and perfusion MRI have important specialized clinical applications, and advanced multi-shell diffusion methods are increasingly used in clinical research, they are not yet uniformly part of routine general-purpose neuroimaging workflows^88,89^. LAMAReg relies on accurate anatomical segmentations to drive registration. In cases where SynthSeg fails, registration quality will likely suffer. While the second stage of registration could alleviate some of these issues, it requires testing in specific disease cases to ensure that it does not disrupt biomarkers of disease. MICAFlow uses Synb0-DISCO to correct dMRI distortions when phase-encoding pairs are unavailable. This approach assumes the availability of standard b=0 images and may underperform relative to reverse-phase encoding corrections on typical data. In rare acquisition schemes or very strong distortions, residual errors may remain. In addition, many of the individual methods and algorithmic components used within our dMRI processing workflow have been independently described and validated in prior studies^44–47^, but full end-to-end validation of the robustness of the combined diffusion pipeline remains an important direction for future work. Our evaluation of MICAFlow on four diverse datasets spanned acquisition sites, parameters, MRI systems, and adult populations. Applications to images of developing and senior brains, as well as atypical MR acquisitions have yet to be demonstrated. Additionally, while epilepsy results in anatomical distortions on MRI which may impact processing^80, 86, 87^, testing of MICAFlow across a wide range of clinical populations is outside of the scope of this evaluation. Although highly optimized, MICAFlow still requires significant processing power for very large or high-resolution datasets. Some steps (*e.g.,* ANTs SyN registration) remain computationally heavy, and due to hardware constraints, we cannot test exhaustively under all conditions, notably the kind of hardware available in lower-resourced clinics (*e.g.,* low RAM, high single-core clock, low clock-multicore, *etc*.). While the runtime was evaluated on commercial hardware, information regarding typical clinician-available hardware is scarce, and likely varies.

## Acknowledgements

IGH received support from the Irma H. Bauer Research Fund. J.D. was supported by a Natural Science and Engineering Research Council of Canada Post Doctoral Fellowship award (NSERC-PDF), Helmholtz International BigBrain Analytics and Learning Laboratory (HIBALL), Healthy Brains and Healthy Lives (HBHL), the Centre for Aging and Brain Health Innovation (CABHI), and the Molson Engineering fellowship of the Montreal Neurological Institute. P.B was funded by FRQ-NT. D.M. is supported by a scholarship from the Fonds de recherche du Québec (award 373207). D.G.C is funded by the FRQ-S and the Savoy Foundation. E.S. acknowledges funding from the Vanier Canada Graduate Scholarship. A.N. was funded by CIHR and FRQ-S. M.S. received support from CIHR. K.X. is supported by the China Scholarship Council and the Savoy Foundation. JL is funded by the Fonds de la Recherche du Québec – Santé (FRQS) and the Ministère de la Santé et des Services sociaux du Québec (MSSS). Y.H. is funded by the Helmholtz International BigBrain Analytics and Learning Laboratory (HIBALL) and the Quebec BioImaging Network (QBIN). J.C. was funded by the Vanier Canada Graduate Scholarship. R.D and G.Z. were funded by the China Scholarship Council. F.F. received support from the Savoy Foundation. RRC received support from the Fonds de la Recherche du Québec – Santé (FRQ-S), the Montreal Neurological Institute Jeanne Timmins Costello Fellowship, and the Healthy Brains, Healthy Lives – Entrepreneur Postdoc Fellowship. BCB acknowledges support from the Canadian Institutes of Health Research, CIHR (FDN-154298, PJT-174995), SickKids Foundation (NI17-039), Natural Sciences and Engineering Research Council (NSERC; Discovery-1304413), Azrieli Center for Autism Research of the Montreal Neurological Institute (ACAR), BrainCanada, FRQ-S, the Helmholtz International BigBrain Analytics and Learning Laboratory (Hiball), the Canada Research Chairs Program (CRC), and the Centre of Excellence in Epilepsy at the Neuro (CEEN).

## Conflict of interest

BCB, JDK, IGH, RRC, and JC hold stock in BrainScores Inc.

