## Supplementary Tables for "MICAFlow: Fast and robust MRI preprocessing bridging research neuroimaging and clinical practice"

**Supplementary Table 1**

| **CPU** | AMD Ryzen 9 7900X |
| --- | --- |
| **Single Core Clock Speed** | 4.7 GHz (5.6 GHz Boost) |
| **Number of cores** | 12 core (24 thread) |
| **RAM** | 64 GB |
| **RAM Speed** | 4800 MHz |
| **Hard Drive** | WD_BLACK SN750 |
| **Hard Drive Read/Write** | Up to 3600 MB/s R, 2900 MB/s W |

**Supplementary Table 2**

| **Registration** | **Metric** | **Comp** | **X̄ Ref** | **X̄ Comp** | **T_stat** | **P_Raw** | **P_Adj** |
| --- | --- | --- | --- | --- | --- | --- | --- |
| T1w/ T1w | MI | ANTs | -0.52201 | -0.52124 | -1.06158 | 0.289 | 0.289 |
| T1w/T1w | MI | EasyReg | -0.52201 | -0.46908 | -12.5824 | 1.779e-29 | 3.557e-29 |
| T1w/T1w | MIND | ANTs | 0.009117 | 0.00917 | -3.10943 | 0.002 | 0.003 |
| T1w/T1w | MIND | EasyReg | 0.009117 | 0.01055 | -14.1622 | 2.880e-35 | 1.728e-34 |
| T1w/T1w | NGF | ANTs | -0.04654 | -0.04637 | -1.95072 | 0.052 | 0.062 |
| T1w/T1w | NGF | EasyReg | -0.04654 | -0.04045 | -13.254 | 6.447e-32 | 1.934e-31 |
| FLAIR/T1w | MI | ANTs | -0.44022 | -0.4401 | -0.73099 | 0.466 | 0.560 |
| FLAIR/T1w | MI | EasyReg | -0.44022 | -0.42769 | -49.4597 | 4.694e-75 | 2.816e-74 |
| FLAIR/T1w | MIND | ANTs | 0.010494 | 0.01059 | -22.3731 | 6.000e-42 | 1.200e-41 |
| FLAIR/T1w | MIND | EasyReg | 0.010494 | 0.01049 | 0.3602 | 0.719 | 0.719 |
| FLAIR/T1w | NGF | ANTs | -0.03964 | -0.03919 | -25.6597 | 2.784e-47 | 8.351e-47 |
| FLAIR/T1w | NGF | EasyReg | -0.03964 | -0.03958 | -2.34862 | 0.021 | 0.031 |
| dMRI/T1w | MI | ANTs | -0.38967 | -0.37782 | -19.7499 | 6.195e-54 | 9.292e-54 |
| dMRI/T1w | MI | FSL | -0.38967 | -0.36093 | -30.4175 | 2.896e-88 | 2.606e-87 |
| dMRI/T1w | MI | EasyReg | -0.38967 | -0.37622 | -12.4201 | 3.354e-28 | 3.354e-28 |
| dMRI/T1w | MIND | ANTs | 0.01180 | 0.01229 | -20.9608 | 4.137e-58 | 7.447e-58 |
| dMRI/T1w | MIND | FSL | 0.01180 | 0.01238 | -24.9055 | 2.943e-71 | 8.829e-71 |
| dMRI/T1w | MIND | EasyReg | 0.01180 | 0.01213 | -12.5759 | 9.789e-29 | 1.101e-28 |
| dMRI/T1w | NGF | ANTs | -0.03596 | -0.03395 | -23.033 | 4.127e-65 | 9.285e-65 |
| dMRI/T1w | NGF | FSL | -0.03596 | -0.03304 | -27.1452 | 2.286e-78 | 1.029e-77 |
| dMRI/T1w | NGF | EasyReg | -0.03596 | -0.0339 | -14.4172 | 3.764e-35 | 4.840e-35 |

**Supplementary Table 3**

| **Modality** | **Aug** | **Strength** | **Metric** | **Method** | **X̄ Lamareg** | **X̄ Ref** | **X̄ Diff** | **P_raw** | **P_adj** |
| --- | --- | --- | --- | --- | --- | --- | --- | --- | --- |
| T1w | Noise | Baseline | MI | Ants (A) | -0.52201 | -0.52124 | -0.00077 | 0.289273 | 0.289273 |
| T1w | Noise | Baseline | MI | Easyreg (E) | -0.52201 | -0.46908 | -0.05293 | 1.78e-29 | 3.56e-29 |
| T1w | Noise | 2.50% | MI | Ants (A) | -0.52279 | -0.52188 | -0.00091 | 2.24e-45 | 4.47e-45 |
| T1w | Noise | 2.50% | MI | Easyreg (E) | -0.52457 | -0.48142 | -0.04315 | 1.05e-16 | 1.05e-16 |
| T1w | Noise | 5% | MI | Ants (A) | -0.52589 | -0.52485 | -0.00104 | 8.87e-09 | 8.87e-09 |
| T1w | Noise | 5% | MI | Easyreg (E) | -0.52599 | -0.48222 | -0.04376 | 5.11e-23 | 1.02e-22 |
| T1w | Noise | 10% | MI | Ants (A) | -0.44364 | -0.52024 | 0.0766 | 1.76e-16 | 3.51e-16 |
| T1w | Noise | 10% | MI | Easyreg (E) | -0.44313 | -0.47226 | 0.029128 | 2.62e-05 | 2.62e-05 |
| T1w | Motion | Baseline | MI | Ants (A) | -0.52201 | -0.52124 | -0.00077 | 0.289273 | 0.289273 |
| T1w | Motion | Baseline | MI | Easyreg (E) | -0.52201 | -0.46908 | -0.05293 | 1.78e-29 | 3.56e-29 |
| T1w | Motion | 5deg | MI | Ants (A) | -0.49836 | -0.49936 | 0.000996 | 9.94e-06 | 9.94e-06 |
| T1w | Motion | 5deg | MI | Easyreg (E) | -0.49836 | -0.49112 | -0.00724 | 9.63e-44 | 1.93e-43 |
| T1w | Motion | 10deg | MI | Ants (A) | -0.50324 | -0.50417 | 0.00093 | 1.29e-07 | 1.29e-07 |
| T1w | Motion | 10deg | MI | Easyreg (E) | -0.50324 | -0.49233 | -0.0109 | 3.61e-10 | 7.21e-10 |
| T1w | Motion | 20deg | MI | Ants (A) | -0.50325 | -0.50396 | 0.000706 | 0.01905 | 0.01905 |
| T1w | Motion | 20deg | MI | Easyreg (E) | -0.50325 | -0.47723 | -0.02602 | 1.60e-18 | 3.19e-18 |
| Flair | Noise | Baseline | MI | Ants (A) | -0.44022 | -0.4401 | -0.00012 | 0.466396 | 0.466396 |
| Flair | Noise | Baseline | MI | Easyreg (E) | -0.44022 | -0.42769 | -0.01253 | 4.69e-75 | 9.39e-75 |
| Flair | Noise | 2.50% | MI | Ants (A) | -0.43973 | -0.43912 | -0.0006 | 0.000314 | 0.000314 |
| Flair | Noise | 2.50% | MI | Easyreg (E) | -0.43973 | -0.42766 | -0.01207 | 7.83e-74 | 1.57e-73 |
| Flair | Noise | 5% | MI | Ants (A) | -0.43931 | -0.43855 | -0.00076 | 4.34e-05 | 4.34e-05 |
| Flair | Noise | 5% | MI | Easyreg (E) | -0.43931 | -0.40711 | -0.0322 | 1.93e-12 | 3.86e-12 |
| Flair | Noise | 10% | MI | Ants (A) | -0.42505 | -0.43749 | 0.012447 | 0.054576 | 0.054576 |
| Flair | Noise | 10% | MI | Easyreg (E) | -0.42505 | -0.36862 | -0.05642 | 7.29e-12 | 1.46e-11 |
| Flair | Motion | Baseline | MI | Ants (A) | -0.44022 | -0.4401 | -0.00012 | 0.466396 | 0.466396 |
| Flair | Motion | Baseline | MI | Easyreg (E) | -0.44022 | -0.42769 | -0.01253 | 4.69e-75 | 9.39e-75 |
| Flair | Motion | 5deg | MI | Ants (A) | -0.42845 | -0.42839 | -6.34e-05 | 0.784845 | 0.784845 |
| Flair | Motion | 5deg | MI | Easyreg (E) | -0.42845 | -0.42027 | -0.00819 | 7.99e-49 | 1.60e-48 |
| Flair | Motion | 10deg | MI | Ants (A) | -0.42606 | -0.42629 | 0.000233 | 0.674103 | 0.674103 |
| Flair | Motion | 10deg | MI | Easyreg (E) | -0.42606 | -0.41848 | -0.00757 | 1.76e-29 | 3.51e-29 |
| Flair | Motion | 20deg | MI | Ants (A) | -0.42771 | -0.42769 | -2.43e-05 | 0.974687 | 0.974687 |
| Flair | Motion | 20deg | MI | Easyreg (E) | -0.42771 | -0.41741 | -0.0103 | 1.28e-27 | 2.57e-27 |
| Dmri | Noise | Baseline | MI | Ants (A) | -0.38967 | -0.37782 | -0.01184 | 6.19e-54 | 9.29e-54 |
| Dmri | Noise | Baseline | MI | Easyreg (E) | -0.38967 | -0.37622 | -0.01345 | 3.35e-28 | 3.35e-28 |
| Dmri | Noise | Baseline | MI | Fsl (F) | -0.38967 | -0.36093 | -0.02873 | 2.90e-88 | 8.69e-88 |
| Dmri | Noise | 2.50% | MI | Ants (A) | -0.39045 | -0.37698 | -0.01346 | 6.21e-61 | 9.32e-61 |
| Dmri | Noise | 2.50% | MI | Easyreg (E) | -0.39022 | -0.38025 | -0.00998 | 2.67e-30 | 2.67e-30 |
| Dmri | Noise | 2.50% | MI | Fsl (F) | -0.39045 | -0.36104 | -0.02941 | 1.45e-88 | 4.35e-88 |
| Dmri | Noise | 5% | MI | Ants (A) | -0.39075 | -0.3789 | -0.01185 | 2.80e-55 | 4.20e-55 |
| Dmri | Noise | 5% | MI | Easyreg (E) | -0.39075 | -0.38176 | -0.00899 | 4.99e-52 | 4.99e-52 |
| Dmri | Noise | 5% | MI | Fsl (F) | -0.39075 | -0.36088 | -0.02987 | 2.02e-88 | 6.06e-88 |
| Dmri | Noise | 10% | MI | Ants (A) | -0.3901 | -0.37946 | -0.01065 | 2.72e-47 | 2.72e-47 |
| Dmri | Noise | 10% | MI | Easyreg (E) | -0.3901 | -0.38196 | -0.00814 | 2.60e-48 | 3.90e-48 |
| Dmri | Noise | 10% | MI | Fsl (F) | -0.3901 | -0.3609 | -0.02921 | 3.70e-84 | 1.11e-83 |
| Dmri | Motion | Baseline | MI | Ants (A) | -0.38967 | -0.37782 | -0.01184 | 6.19e-54 | 9.29e-54 |
| Dmri | Motion | Baseline | MI | Easyreg (E) | -0.38967 | -0.37622 | -0.01345 | 3.35e-28 | 3.35e-28 |
| Dmri | Motion | Baseline | MI | Fsl (F) | -0.38967 | -0.36093 | -0.02873 | 2.90e-88 | 8.69e-88 |
| Dmri | Motion | 5deg | MI | Ants (A) | -0.37932 | -0.36803 | -0.01129 | 6.72e-37 | 1.01e-36 |
| Dmri | Motion | 5deg | MI | Easyreg (E) | -0.37897 | -0.37291 | -0.00606 | 1.15e-06 | 1.15e-06 |
| Dmri | Motion | 5deg | MI | Fsl (F) | -0.37932 | -0.35892 | -0.0204 | 1.81e-52 | 5.42e-52 |
| Dmri | Motion | 10deg | MI | Ants (A) | -0.3799 | -0.36788 | -0.01202 | 1.64e-46 | 4.93e-46 |
| Dmri | Motion | 10deg | MI | Easyreg (E) | -0.3799 | -0.37539 | -0.00451 | 0.006661 | 0.006661 |
| Dmri | Motion | 10deg | MI | Fsl (F) | -0.3799 | -0.36004 | -0.01986 | 9.76e-44 | 1.46e-43 |
| Dmri | Motion | 20deg | MI | Ants (A) | -0.37872 | -0.36563 | -0.01309 | 2.21e-33 | 6.62e-33 |
| Dmri | Motion | 20deg | MI | Easyreg (E) | -0.37804 | -0.3713 | -0.00674 | 0.000548 | 0.000548 |
| Dmri | Motion | 20deg | MI | Fsl (F) | -0.37872 | -0.36096 | -0.01776 | 2.02e-31 | 3.03e-31 |
| T1w | Noise | Baseline | Mind | Ants (A) | 0.009117 | 0.009169 | -5.22e-05 | 0.002053 | 0.002053 |
| T1w | Noise | Baseline | Mind | Easyreg (E) | 0.009117 | 0.010552 | -0.00143 | 2.88e-35 | 5.76e-35 |
| T1w | Noise | 2.50% | Mind | Ants (A) | 0.009119 | 0.009173 | -5.44e-05 | 3.43e-64 | 6.87e-64 |
| T1w | Noise | 2.50% | Mind | Easyreg (E) | 0.009231 | 0.010451 | -0.00122 | 3.08e-21 | 3.08e-21 |
| T1w | Noise | 5% | Mind | Ants (A) | 0.00938 | 0.009436 | -5.59e-05 | 3.07e-30 | 6.14e-30 |
| T1w | Noise | 5% | Mind | Easyreg (E) | 0.009381 | 0.010865 | -0.00148 | 3.25e-27 | 3.25e-27 |
| T1w | Noise | 10% | Mind | Ants (A) | 0.010479 | 0.009258 | 0.001221 | 6.03e-17 | 1.21e-16 |
| T1w | Noise | 10% | Mind | Easyreg (E) | 0.010479 | 0.011065 | -0.00059 | 3.94e-10 | 3.94e-10 |
| T1w | Motion | Baseline | Mind | Ants (A) | 0.009117 | 0.009169 | -5.22e-05 | 0.002053 | 0.002053 |
| T1w | Motion | Baseline | Mind | Easyreg (E) | 0.009117 | 0.010552 | -0.00143 | 2.88e-35 | 5.76e-35 |
| T1w | Motion | 5deg | Mind | Ants (A) | 0.009817 | 0.009784 | 3.30e-05 | 3.96e-05 | 3.96e-05 |
| T1w | Motion | 5deg | Mind | Easyreg (E) | 0.009817 | 0.010023 | -0.00021 | 8.11e-27 | 1.62e-26 |
| T1w | Motion | 10deg | Mind | Ants (A) | 0.009917 | 0.00989 | 2.68e-05 | 2.25e-06 | 2.25e-06 |
| T1w | Motion | 10deg | Mind | Easyreg (E) | 0.009917 | 0.010145 | -0.00023 | 8.31e-10 | 1.66e-09 |
| T1w | Motion | 20deg | Mind | Ants (A) | 0.009908 | 0.009887 | 2.10e-05 | 0.008416 | 0.008416 |
| T1w | Motion | 20deg | Mind | Easyreg (E) | 0.009908 | 0.01047 | -0.00056 | 2.28e-16 | 4.56e-16 |
| Flair | Noise | Baseline | Mind | Ants (A) | 0.010494 | 0.010592 | -9.78e-05 | 6.00e-42 | 1.20e-41 |
| Flair | Noise | Baseline | Mind | Easyreg (E) | 0.010494 | 0.010491 | 3.44e-06 | 0.7194 | 0.7194 |
| Flair | Noise | 2.50% | Mind | Ants (A) | 0.010509 | 0.010596 | -8.77e-05 | 1.15e-45 | 2.29e-45 |
| Flair | Noise | 2.50% | Mind | Easyreg (E) | 0.010509 | 0.010487 | 2.12e-05 | 0.028987 | 0.028987 |
| Flair | Noise | 5% | Mind | Ants (A) | 0.01052 | 0.010618 | -9.83e-05 | 1.95e-45 | 3.91e-45 |
| Flair | Noise | 5% | Mind | Easyreg (E) | 0.01052 | 0.011095 | -0.00057 | 1.68e-13 | 1.68e-13 |
| Flair | Noise | 10% | Mind | Ants (A) | 0.010622 | 0.010655 | -3.32e-05 | 0.243282 | 0.243282 |
| Flair | Noise | 10% | Mind | Easyreg (E) | 0.010622 | 0.012317 | -0.00169 | 2.21e-30 | 4.42e-30 |
| Flair | Motion | Baseline | Mind | Ants (A) | 0.010494 | 0.010592 | -9.78e-05 | 6.00e-42 | 1.20e-41 |
| Flair | Motion | Baseline | Mind | Easyreg (E) | 0.010494 | 0.010491 | 3.44e-06 | 0.7194 | 0.7194 |
| Flair | Motion | 5deg | Mind | Ants (A) | 0.010903 | 0.010976 | -7.36e-05 | 2.90e-11 | 5.81e-11 |
| Flair | Motion | 5deg | Mind | Easyreg (E) | 0.010903 | 0.010849 | 5.33e-05 | 0.002744 | 0.002744 |
| Flair | Motion | 10deg | Mind | Ants (A) | 0.010971 | 0.011012 | -4.12e-05 | 0.008163 | 0.008163 |
| Flair | Motion | 10deg | Mind | Easyreg (E) | 0.010971 | 0.010909 | 6.22e-05 | 0.007616 | 0.008163 |
| Flair | Motion | 20deg | Mind | Ants (A) | 0.010918 | 0.010992 | -7.44e-05 | 0.020627 | 0.041253 |
| Flair | Motion | 20deg | Mind | Easyreg (E) | 0.010918 | 0.010944 | -2.61e-05 | 0.475827 | 0.475827 |
| Dmri | Noise | Baseline | Mind | Ants (A) | 0.011803 | 0.012293 | -0.00049 | 4.14e-58 | 6.21e-58 |
| Dmri | Noise | Baseline | Mind | Easyreg (E) | 0.011803 | 0.012137 | -0.00033 | 9.79e-29 | 9.79e-29 |
| Dmri | Noise | Baseline | Mind | Fsl (F) | 0.011803 | 0.012375 | -0.00057 | 2.94e-71 | 8.83e-71 |
| Dmri | Noise | 2.50% | Mind | Ants (A) | 0.011807 | 0.012287 | -0.00048 | 1.34e-54 | 2.01e-54 |
| Dmri | Noise | 2.50% | Mind | Easyreg (E) | 0.011797 | 0.011984 | -0.00019 | 1.64e-19 | 1.64e-19 |
| Dmri | Noise | 2.50% | Mind | Fsl (F) | 0.011807 | 0.012375 | -0.00057 | 4.39e-70 | 1.32e-69 |
| Dmri | Noise | 5% | Mind | Ants (A) | 0.011822 | 0.0123 | -0.00048 | 1.50e-55 | 2.26e-55 |
| Dmri | Noise | 5% | Mind | Easyreg (E) | 0.011822 | 0.011933 | -0.00011 | 1.98e-31 | 1.98e-31 |
| Dmri | Noise | 5% | Mind | Fsl (F) | 0.011822 | 0.012375 | -0.00055 | 3.01e-67 | 9.04e-67 |
| Dmri | Noise | 10% | Mind | Ants (A) | 0.011887 | 0.012354 | -0.00047 | 1.00e-54 | 1.50e-54 |
| Dmri | Noise | 10% | Mind | Easyreg (E) | 0.011887 | 0.011959 | -7.14e-05 | 3.54e-12 | 3.54e-12 |
| Dmri | Noise | 10% | Mind | Fsl (F) | 0.011887 | 0.012377 | -0.00049 | 5.55e-57 | 1.67e-56 |
| Dmri | Motion | Baseline | Mind | Ants (A) | 0.011803 | 0.012293 | -0.00049 | 4.14e-58 | 6.21e-58 |
| Dmri | Motion | Baseline | Mind | Easyreg (E) | 0.011803 | 0.012137 | -0.00033 | 9.79e-29 | 9.79e-29 |
| Dmri | Motion | Baseline | Mind | Fsl (F) | 0.011803 | 0.012375 | -0.00057 | 2.94e-71 | 8.83e-71 |
| Dmri | Motion | 5deg | Mind | Ants (A) | 0.012189 | 0.012673 | -0.00048 | 4.30e-64 | 1.29e-63 |
| Dmri | Motion | 5deg | Mind | Easyreg (E) | 0.012191 | 0.01236 | -0.00017 | 1.92e-11 | 1.92e-11 |
| Dmri | Motion | 5deg | Mind | Fsl (F) | 0.012189 | 0.012485 | -0.0003 | 4.96e-28 | 7.45e-28 |
| Dmri | Motion | 10deg | Mind | Ants (A) | 0.012191 | 0.012706 | -0.00052 | 1.11e-63 | 3.33e-63 |
| Dmri | Motion | 10deg | Mind | Easyreg (E) | 0.012191 | 0.012396 | -0.0002 | 2.34e-14 | 3.50e-14 |
| Dmri | Motion | 10deg | Mind | Fsl (F) | 0.012191 | 0.012415 | -0.00022 | 5.11e-14 | 5.11e-14 |
| Dmri | Motion | 20deg | Mind | Ants (A) | 0.012204 | 0.01271 | -0.00051 | 2.72e-59 | 8.17e-59 |
| Dmri | Motion | 20deg | Mind | Easyreg (E) | 0.012173 | 0.012515 | -0.00034 | 1.42e-28 | 2.13e-28 |
| Dmri | Motion | 20deg | Mind | Fsl (F) | 0.012204 | 0.012353 | -0.00015 | 1.67e-06 | 1.67e-06 |
| T1w | Noise | Baseline | Ngf | Ants (A) | -0.04654 | -0.04637 | -0.00017 | 0.052015 | 0.052015 |
| T1w | Noise | Baseline | Ngf | Easyreg (E) | -0.04654 | -0.04045 | -0.00609 | 6.45e-32 | 1.29e-31 |
| T1w | Noise | 2.50% | Ngf | Ants (A) | -0.04659 | -0.04642 | -0.00017 | 2.45e-60 | 4.89e-60 |
| T1w | Noise | 2.50% | Ngf | Easyreg (E) | -0.04648 | -0.0411 | -0.00538 | 2.18e-18 | 2.18e-18 |
| T1w | Noise | 5% | Ngf | Ants (A) | -0.04624 | -0.04605 | -0.00019 | 4.93e-17 | 4.93e-17 |
| T1w | Noise | 5% | Ngf | Easyreg (E) | -0.04625 | -0.04067 | -0.00559 | 1.68e-26 | 3.37e-26 |
| T1w | Noise | 10% | Ngf | Ants (A) | -0.03818 | -0.04616 | 0.007971 | 6.86e-17 | 1.37e-16 |
| T1w | Noise | 10% | Ngf | Easyreg (E) | -0.03819 | -0.03936 | 0.001174 | 0.057772 | 0.057772 |
| T1w | Motion | Baseline | Ngf | Ants (A) | -0.04654 | -0.04637 | -0.00017 | 0.052015 | 0.052015 |
| T1w | Motion | Baseline | Ngf | Easyreg (E) | -0.04654 | -0.04045 | -0.00609 | 6.45e-32 | 1.29e-31 |
| T1w | Motion | 5deg | Ngf | Ants (A) | -0.04388 | -0.04397 | 8.45e-05 | 0.02589 | 0.02589 |
| T1w | Motion | 5deg | Ngf | Easyreg (E) | -0.04388 | -0.04305 | -0.00083 | 6.00e-36 | 1.20e-35 |
| T1w | Motion | 10deg | Ngf | Ants (A) | -0.0441 | -0.04416 | 5.60e-05 | 0.02572 | 0.02572 |
| T1w | Motion | 10deg | Ngf | Easyreg (E) | -0.0441 | -0.04296 | -0.00114 | 4.80e-11 | 9.60e-11 |
| T1w | Motion | 20deg | Ngf | Ants (A) | -0.04415 | -0.04417 | 2.69e-05 | 0.503604 | 0.503604 |
| T1w | Motion | 20deg | Ngf | Easyreg (E) | -0.04415 | -0.04133 | -0.00281 | 3.75e-19 | 7.51e-19 |
| Flair | Noise | Baseline | Ngf | Ants (A) | -0.03964 | -0.03919 | -0.00045 | 2.78e-47 | 5.57e-47 |
| Flair | Noise | Baseline | Ngf | Easyreg (E) | -0.03964 | -0.03958 | -6.49e-05 | 0.020698 | 0.020698 |
| Flair | Noise | 2.50% | Ngf | Ants (A) | -0.03958 | -0.03917 | -0.00041 | 2.07e-51 | 4.15e-51 |
| Flair | Noise | 2.50% | Ngf | Easyreg (E) | -0.03958 | -0.03958 | -2.81e-06 | 0.922233 | 0.922233 |
| Flair | Noise | 5% | Ngf | Ants (A) | -0.03953 | -0.03908 | -0.00045 | 1.92e-50 | 3.85e-50 |
| Flair | Noise | 5% | Ngf | Easyreg (E) | -0.03953 | -0.03681 | -0.00272 | 1.66e-11 | 1.66e-11 |
| Flair | Noise | 10% | Ngf | Ants (A) | -0.0385 | -0.03893 | 0.000424 | 0.342254 | 0.342254 |
| Flair | Noise | 10% | Ngf | Easyreg (E) | -0.0385 | -0.0309 | -0.0076 | 1.06e-21 | 2.12e-21 |
| Flair | Motion | Baseline | Ngf | Ants (A) | -0.03964 | -0.03919 | -0.00045 | 2.78e-47 | 5.57e-47 |
| Flair | Motion | Baseline | Ngf | Easyreg (E) | -0.03964 | -0.03958 | -6.49e-05 | 0.020698 | 0.020698 |
| Flair | Motion | 5deg | Ngf | Ants (A) | -0.03814 | -0.03774 | -0.00041 | 6.97e-14 | 1.39e-13 |
| Flair | Motion | 5deg | Ngf | Easyreg (E) | -0.03814 | -0.03823 | 8.44e-05 | 0.164747 | 0.164747 |
| Flair | Motion | 10deg | Ngf | Ants (A) | -0.03781 | -0.03758 | -0.00023 | 0.005352 | 0.010703 |
| Flair | Motion | 10deg | Ngf | Easyreg (E) | -0.03781 | -0.03791 | 9.92e-05 | 0.284174 | 0.284174 |
| Flair | Motion | 20deg | Ngf | Ants (A) | -0.03804 | -0.03767 | -0.00037 | 0.021805 | 0.032158 |
| Flair | Motion | 20deg | Ngf | Easyreg (E) | -0.03804 | -0.03768 | -0.00036 | 0.032158 | 0.032158 |
| Dmri | Noise | Baseline | Ngf | Ants (A) | -0.03596 | -0.03395 | -0.00201 | 4.13e-65 | 6.19e-65 |
| Dmri | Noise | Baseline | Ngf | Easyreg (E) | -0.03596 | -0.0339 | -0.00206 | 3.76e-35 | 3.76e-35 |
| Dmri | Noise | Baseline | Ngf | Fsl (F) | -0.03596 | -0.03304 | -0.00292 | 2.29e-78 | 6.86e-78 |
| Dmri | Noise | 2.50% | Ngf | Ants (A) | -0.03593 | -0.03386 | -0.00207 | 1.28e-62 | 1.92e-62 |
| Dmri | Noise | 2.50% | Ngf | Easyreg (E) | -0.03596 | -0.03471 | -0.00125 | 2.02e-29 | 2.02e-29 |
| Dmri | Noise | 2.50% | Ngf | Fsl (F) | -0.03593 | -0.03304 | -0.00288 | 2.86e-76 | 8.57e-76 |
| Dmri | Noise | 5% | Ngf | Ants (A) | -0.03585 | -0.03382 | -0.00203 | 4.78e-62 | 4.78e-62 |
| Dmri | Noise | 5% | Ngf | Easyreg (E) | -0.03585 | -0.03501 | -0.00084 | 3.76e-67 | 5.64e-67 |
| Dmri | Noise | 5% | Ngf | Fsl (F) | -0.03585 | -0.03305 | -0.0028 | 1.44e-72 | 4.33e-72 |
| Dmri | Noise | 10% | Ngf | Ants (A) | -0.03555 | -0.03357 | -0.00198 | 8.22e-60 | 1.23e-59 |
| Dmri | Noise | 10% | Ngf | Easyreg (E) | -0.03555 | -0.03488 | -0.00067 | 7.52e-38 | 7.52e-38 |
| Dmri | Noise | 10% | Ngf | Fsl (F) | -0.03555 | -0.03304 | -0.0025 | 7.88e-62 | 2.36e-61 |
| Dmri | Motion | Baseline | Ngf | Ants (A) | -0.03596 | -0.03395 | -0.00201 | 4.13e-65 | 6.19e-65 |
| Dmri | Motion | Baseline | Ngf | Easyreg (E) | -0.03596 | -0.0339 | -0.00206 | 3.76e-35 | 3.76e-35 |
| Dmri | Motion | Baseline | Ngf | Fsl (F) | -0.03596 | -0.03304 | -0.00292 | 2.29e-78 | 6.86e-78 |
| Dmri | Motion | 5deg | Ngf | Ants (A) | -0.03396 | -0.03204 | -0.00191 | 1.58e-60 | 4.75e-60 |
| Dmri | Motion | 5deg | Ngf | Easyreg (E) | -0.03395 | -0.03276 | -0.00119 | 1.09e-17 | 1.09e-17 |
| Dmri | Motion | 5deg | Ngf | Fsl (F) | -0.03396 | -0.03251 | -0.00145 | 2.12e-29 | 3.17e-29 |
| Dmri | Motion | 10deg | Ngf | Ants (A) | -0.03385 | -0.03181 | -0.00203 | 5.46e-53 | 1.64e-52 |
| Dmri | Motion | 10deg | Ngf | Easyreg (E) | -0.03385 | -0.03246 | -0.00138 | 5.71e-21 | 8.56e-21 |
| Dmri | Motion | 10deg | Ngf | Fsl (F) | -0.03385 | -0.03283 | -0.00102 | 1.01e-11 | 1.01e-11 |
| Dmri | Motion | 20deg | Ngf | Ants (A) | -0.03357 | -0.03151 | -0.00205 | 3.45e-43 | 1.03e-42 |
| Dmri | Motion | 20deg | Ngf | Easyreg (E) | -0.03358 | -0.03111 | -0.00247 | 3.49e-41 | 5.23e-41 |
| Dmri | Motion | 20deg | Ngf | Fsl (F) | -0.03357 | -0.03297 | -0.0006 | 0.000274 | 0.000274 |

**Supplementary Table 4**

| **Dataset** | **Category** | **Mean** | **Std** | **Mean_Intra** | **Mean_Inter** |
| --- | --- | --- | --- | --- | --- |
| MICs | Linear | 51.708 | 2.485 | 0.933 | 0.416 |
| MICs | Nonlinear | 36.955 | 1.825 | 0.937 | 0.567 |
| EpiC | Linear | 40.828 | 2.463 | 0.890 | 0.482 |
| EpiC | Nonlinear | 27.975 | 1.661 | 0.902 | 0.623 |
| PNI | Linear | 39.020 | 4.449 | 0.777 | 0.387 |
| PNI | Nonlinear | 31.460 | 4.297 | 0.799 | 0.484 |
| AmTrT | Linear | 39.601 | 3.621 | 0.912 | 0.516 |
| AmTrT | Nonlinear | 24.841 | 4.009 | 0.896 | 0.648 |

**Supplementary Table 5**

| **Dataset** | **Category** | **Metric** | **Mean** | **Std** |
| --- | --- | --- | --- | --- |
| MICs | T1_Linear | MI | -5.65E-01 | 4.10E-03 |
| MICs | T1_SyN | MI | -6.00E-01 | 4.34E-03 |
| MICs | FLAIR | MI | -4.29E-01 | 2.75E-02 |
| MICs | dMRI | MI | -4.01E-01 | 2.49E-02 |
| MICs | T1_Linear | MIND | 2.04E-02 | 1.96E-04 |
| MICs | T1_SyN | MIND | 1.92E-02 | 2.10E-04 |
| MICs | FLAIR | MIND | 1.10E-02 | 1.11E-03 |
| MICs | dMRI | MIND | 1.29E-02 | 1.14E-03 |
| MICs | T1_Linear | NGF | -5.31E-02 | 8.24E-04 |
| MICs | T1_SyN | NGF | -5.87E-02 | 8.77E-04 |
| MICs | FLAIR | NGF | -3.89E-02 | 3.12E-03 |
| MICs | dMRI | NGF | -3.24E-02 | 2.86E-03 |
| EpiC | T1_Linear | MI | -5.59E-01 | 1.10E-02 |
| EpiC | T1_SyN | MI | -6.05E-01 | 6.05E-03 |
| EpiC | FLAIR | MI | -4.11E-01 | 3.86E-02 |
| EpiC | dMRI | MI | -3.40E-01 | 3.02E-02 |
| EpiC | T1_Linear | MIND | 2.02E-02 | 3.86E-04 |
| EpiC | T1_SyN | MIND | 1.86E-02 | 4.21E-04 |
| EpiC | FLAIR | MIND | 9.54E-03 | 1.38E-03 |
| EpiC | dMRI | MIND | 1.09E-02 | 1.50E-03 |
| EpiC | T1_Linear | NGF | -5.46E-02 | 1.15E-03 |
| EpiC | T1_SyN | NGF | -6.12E-02 | 1.46E-03 |
| EpiC | FLAIR | NGF | -3.95E-02 | 5.41E-03 |
| EpiC | dMRI | NGF | -2.78E-02 | 3.84E-03 |
| PNI | T1_Linear | MI | -5.64E-01 | 3.00E-03 |
| PNI | T1_SyN | MI | -5.93E-01 | 4.48E-03 |
| PNI | dMRI | MI | -2.84E-01 | 3.40E-02 |
| PNI | T1_Linear | MIND | 2.05E-02 | 2.07E-04 |
| PNI | T1_SyN | MIND | 1.95E-02 | 2.22E-04 |
| PNI | dMRI | MIND | 1.47E-02 | 1.15E-03 |
| PNI | T1_Linear | NGF | -5.19E-02 | 5.73E-04 |
| PNI | T1_SyN | NGF | -5.65E-02 | 8.02E-04 |
| PNI | dMRI | NGF | -2.98E-02 | 2.52E-03 |
| AmTrT | T1_Linear | MI | -5.60E-01 | 1.02E-02 |
| AmTrT | T1_SyN | MI | -6.00E-01 | 8.19E-03 |
| AmTrT | dMRI | MI | -3.27E-01 | 2.71E-02 |
| AmTrT | T1_Linear | MIND | 2.01E-02 | 2.20E-04 |
| AmTrT | T1_SyN | MIND | 1.88E-02 | 2.36E-04 |
| AmTrT | dMRI | MIND | 1.09E-02 | 9.11E-04 |
| AmTrT | T1_Linear | NGF | -5.48E-02 | 9.28E-04 |
| AmTrT | T1_SyN | NGF | -6.14E-02 | 1.11E-03 |
| AmTrT | dMRI | NGF | -3.05E-02 | 3.05E-03 |

**Supplementary Table 6**

| **Dataset** | **Category** | **Mean** | **STD** |
| --- | --- | --- | --- |
| MICs | FA | 0.931173 | 1.61E-02 |
| MICs | ADC | 0.916933 | 2.41E-02 |
| EpiC | FA | 8.67E-01 | 5.40E-02 |
| EpiC | ADC | 0.813109 | 6.47E-02 |
| PNI | FA | 0.643343 | 8.25E-02 |
| PNI | ADC | 0.776989 | 8.27E-02 |
| AmTrT | FA | 0.940533 | 1.79E-02 |
| AmTrT | ADC | 0.849046 | 8.10E-02 |

**Supplementary Table 7**

| **Dataset** | **Source** | **Group** | **Mean** | **STD** |
| --- | --- | --- | --- | --- |
| EpiC | linear | control | 0.890 | 0.033 |
| EpiC | nonlinear | control | 0.902 | 0.025 |
| EpiC | linear | patient | 0.862 | 0.038 |
| EpiC | nonlinear | patient | 0.863 | 0.041 |
| MICs | linear | control | 0.932 | 0.027 |
| MICs | nonlinear | control | 0.935 | 0.013 |
| MICs | linear | patient | 0.889 | 0.054 |
| MICs | nonlinear | patient | 0.913 | 0.037 |

**Supplementary Table 8**

| **Dataset** | **Source** | **Group** | **Identifiability** |
| --- | --- | --- | --- |
| EPIC | linear | patient | 41.243 |
| EPIC | nonlinear | patient | 27.232 |
| EPIC | linear | control | 40.825 |
| EPIC | nonlinear | control | 27.974 |
| MICS | linear | control | 51.460 |
| MICS | nonlinear | control | 36.706 |
| MICS | linear | patient | 48.556 |
| MICS | nonlinear | patient | 35.352 |
